# A biophotoelectrochemical approach to unravelling the role of cyanobacterial cell structures in exoelectrogenesis

**DOI:** 10.1101/2021.04.01.437897

**Authors:** Laura T. Wey, Joshua M. Lawrence, Xiaolong Chen, Robert Clark, David J. Lea-Smith, Jenny Z. Zhang, Christopher J. Howe

## Abstract

Photosynthetic microorganisms can export electrons outside their cells, a phenomenon called exoelectrogenesis, which can be harnessed for solar energy conversion. However, the route electrons take from thylakoid membranes to the cell exterior is not understood. Electrochemistry is a powerful analytical technique for studying electron transfer pathways. Here, we show how photoelectrochemistry can be used to compare electron flux from cyanobacterial cells of different growth stages, species and with the outer layers systematically removed. We show that the periplasmic space contributes significantly to the photocurrent profile complexity of whole cells, indicating that it gates electron transfer in exoelectrogenesis. We found that although components of the type IV pili machinery do not have a role in exoelectrogenesis, they contribute significantly to cell-electrode adherence. This study establishes that analytical photoelectrochemistry and molecular microbiology provide a powerful combination to study exoelectrogenesis, enabling future studies to answer biological questions and advance solar energy conversion applications.

## Introduction

Isolated chloroplasts and photosynthetic microorganisms, including eukaryotic algae and cyanobacteria, can export electrons or reducing equivalents upon illumination in a phenomenon called ‘exoelectrogenesis’.^1, 2^ Exoelectrogenesis is a poorly understood process hypothesised to be involved in response to light stress,^3^ iron acquisition,^4^ or signalling.^5^ Despite our lack of understanding of the mechanism, exoelectrogenesis has been demonstrated to be useful for solar energy conversion applications, for example in biophotovoltaics where exoelectrogenic photosynthetic cells are employed to convert sunlight into electricity.^6–10^ However, gaps in understanding of the mechanism(s) of exoelectrogenesis hinder the development of such novel biotechnologies.

Most studies of the exoelectrogenic activity of cyanobacterial cells have concentrated on the maximum current output under illumination. However, the current output over time under light/dark cycles gives a ‘photocurrent profile’ with complexity (i.e. a pattern of peaks and troughs before reaching a steady-state) that is not seen using isolated photosystem II (**Fig. 1A**), ^8, 11–16^ which is known to be the source of the electrons eventually exported during exoelectrogenesis.^17, 18^ We hypothesised that the complexity in the photocurrent profile results from the route taken by the electrons from photosystem II across the many outer layers of the cyanobacterial cell structure to the electrode **(Fig. 1B).**

**Figure 1.**
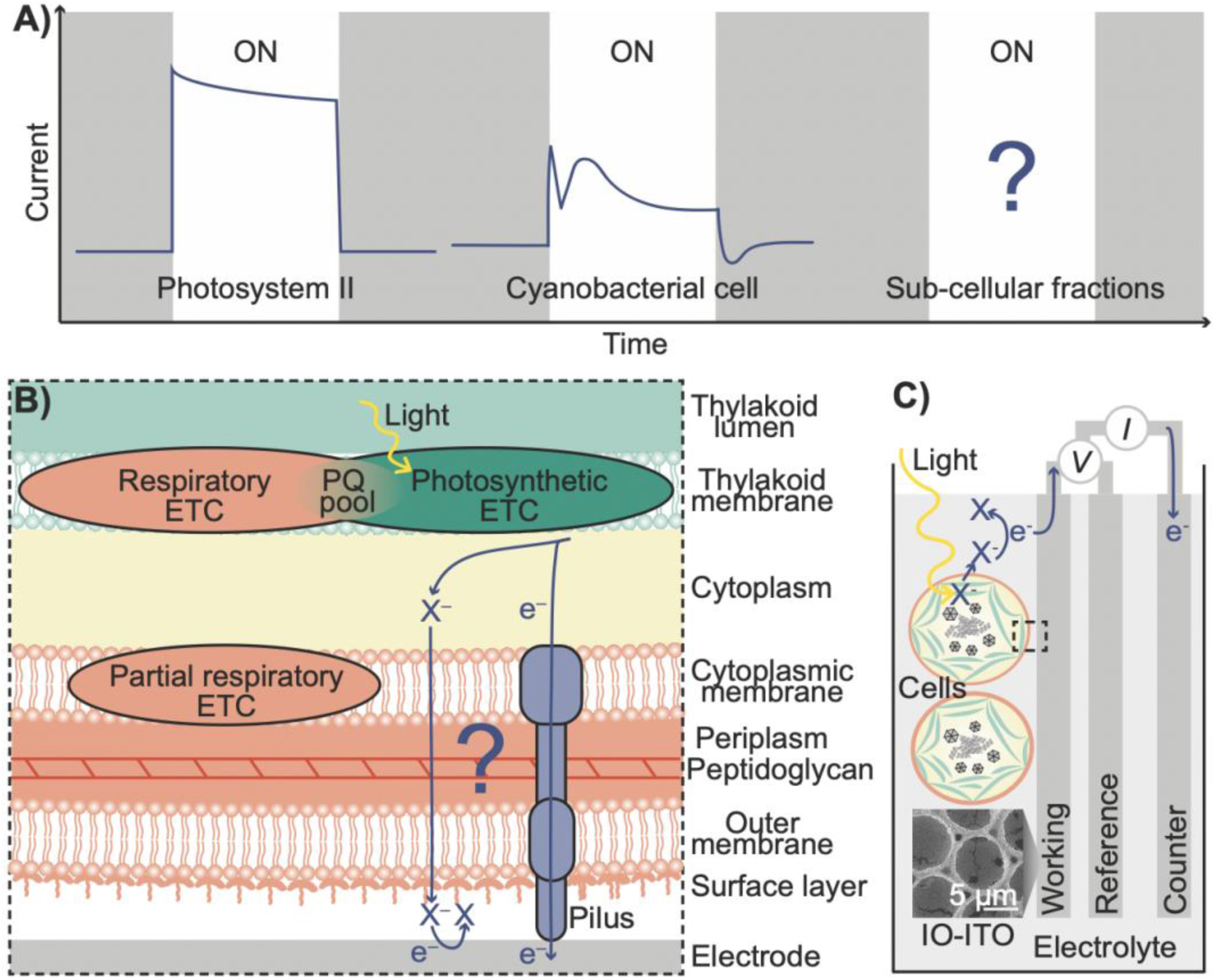
**A)** Schematic representation of chronoamperometric traces under light/dark cycles (i.e. photocurrent profiles) of photosystem II (PSII) protein-films and biofilms of cyanobacterial cells from previous studies.^13^ ON, light on. **B)** Schematic representation of the cell-electrode interface (dashed box from panel C, rotated 90°) showing cyanobacterial cell topology. Postulated electron transfer pathways behind exoelectrogenesis are shown by purple arrows. ‘X’ is an unidentified electron carrier produced and exported by the cells under illumination. More detail of the electron transfer pathways is in Supplementary Figure 1. **C)** Schematic representation of the three-electrode photoelectrochemical set-up used for studying exoelectrogenesis. Cyanobacterial cells or subcellular fractions are interfaced with the working electrode which is referenced to the Ag/AgCl electrode. The panel also shows a scanning electron micrograph of an inverse-opal indium tin oxide (IO-ITO) working electrode used in this study.

The cell structure of Gram-negative cyanobacteria, such as *Synechocystis sp*. PCC6803 (*Synechocystis*), is well characterised. Photosystem II is a part of the photosynthetic electron transport chain, which is located in the thylakoid membranes in cyanobacteria and other oxygenic photosynthetic organisms. The cyanobacterial thylakoid membranes are separated from the cytoplasmic membrane bounding the cell by the cytoplasm.^19^ Outside the cytoplasmic membrane is, collectively referred to as the cell wall, a periplasmic space containing a peptidoglycan layer, an outer membrane, and a surface layer (S-layer) composed of a lattice formed of a single glycoprotein.^20–22^ Extracellular appendages traverse the cytoplasmic membrane and cell wall, including two morphotypes of type IV pili: thick pili, 6-8 nm and more than 2 µm long; and thin pili, 3-4 nm diameter and up to 1 µm long.^23^ There is some evidence of direct extracellular electron transfer via type IV pili,^24, 25^ but recent genetic and conductivity data call this into question.^15^ There is evidence for an endogenous electron mediator (for example, a quinone or NADPH) that enables indirect extracellular electron transfer from the cell to the electrode surface.^13, 26, 27^

The mechanism of electron transfer by isolated proteins, including photosystem II, has been well studied using electrochemistry.^16, 28–31^ Here, we apply photoelectrochemistry to study exoelectrogenesis of cyanobacterial cells **(Fig. 1C**). To identify the cyanobacterial cell structure gating exoelectrogenesis and thus responsible for the photocurrent profile complexity we analysed the photocurrent output of subcellular fractions of *Synechocystis* cells from which different outer layers had been removed. To do this, we first developed a robust microbiological and analytical protocol for comparing the differences in photocurrent profiles of cyanobacterial cells cultured to different growth phases and from different species. We also investigated the effect of cell-electrode adherence on photocurrent outputs, and tested the longevity and efficiency of connection to electrodes (‘wiring’) of subcellular fractions. This top-down biophotoelectrochemical approach has allowed us to identify the importance of the periplasmic space in the gating of cyanobacterial extracellular electron transfer.

## Results

### Photocurrent profile features are conserved in different growth phases and species

To measure the photocurrent profiles of cyanobacteria reproducibly, we first developed a robust microbial and photoelectrochemical protocol. Cells of two widely used model cyanobacterial species *Synechocystis* and *Synechococcus elongatus* PCC7942 (*Synechococcus*) were cultured photoautotrophically in BG11 medium and harvested at specific phases in the growth cycle of cyanobacteria (exponential, early stationary, and late stationary). A concentrated cell suspension with 37.5 nmol of chlorophyll was loaded onto electrodes, which were then left in the dark for 16 h to avoid differences in light exposure immediately before photoelectrochemical analysis. We used state-of-the-art hierarchically structured inverse-opal indium-tin oxide (IO-ITO) electrodes, which boost cell loading and electrochemical signals.^13, 32^ Chronoamperometry was conducted at an applied potential of +0.1 V vs Ag/AgCl (saturated KCl) reference electrode (equivalent to +0.3 V vs Standard Hydrogen Electrode (SHE)), the potential at which the maximum photocurrent is reached for whole *Synechocystis* cells as derived from stepped chronoamperometry.^13^ During chronoamperometry the current was measured each second enabling resolution of the features in the trace, and the loaded electrodes were exposed to cycles of 680 nm light at 50 µmol photons m^−2^ s^−1^ (approximately 1 mW cm^−2^ equivalent) to drive photosynthesis, followed by dark, for 60 s and 90 s respectively (after which periods the steady-state currents in the light and dark had stabilised). The photocurrent was calculated as the difference between the steady-state currents in the light and dark. The photocurrent was normalised to the geometric area of the electrode to obtain photocurrent densities, enabling comparison of outputs between electrodes. In addition, the chlorophyll content of cells remaining on the electrodes after photoelectrochemistry experiments was measured and used as a proxy for the amount of biological material participating in exoelectrogenesis, so different biological samples could be compared.

Comparing the photocurrent outputs of photosynthetic microorganisms between studies is difficult. For example, not all studies report the growth phase, which may affect exoelectrogenesis.^6, 33^ To investigate exoelectrogenesis at different phases in the growth cycle of cyanobacteria, we compared the photoelectrochemistry of cultures of the widely used cyanobacterial species *Synechocystis* at different growth phases: exponential (optical density at 750 nm 0.4), early stationary (OD_750_ = 1), and late stationary (OD_750_ > 1.5) phase.^34^ Under the conditions described above, the cells at the different growth phases gave reproducible photocurrent profiles that were complex (i.e. did not simply reach a steady-state directly) and showed the same features. They all showed two peaks at 5 s and 25 s after illumination, a trough that reached its minimum at 12 s after illumination and a broader trough that reached its minimum at 15 s after the light was turned off (**Fig. 2A, C**). The peaks and troughs in the photocurrent profile were most pronounced for exponential phase cells and the variation in photocurrent profile between biological replicates was smallest for early stationary phase cells. The photocurrent outputs of the different cells loaded were determined, with exponential phase cells yielding a photocurrent density (0.21 ± 0.03 µA cm^−2^) approximately double that from early stationary phase cells (0.110 ± 0.004 µA cm^−2^) (**Fig. 2D**). When normalised to the chlorophyll content, the photocurrent densities of electrodes loaded with cells from exponential, early stationary and late stationary phases were similar (early stationary phase: 160 ± 74 A cm^−2^ mol^−1^(Chl a)) (**Fig. 2E**). For subsequent analytical photoelectrochemical experiments, cells harvested at early stationary phase (OD_750_ = 1) were used, as they displayed the least variability between replicate measurements.

**Figure 2.**
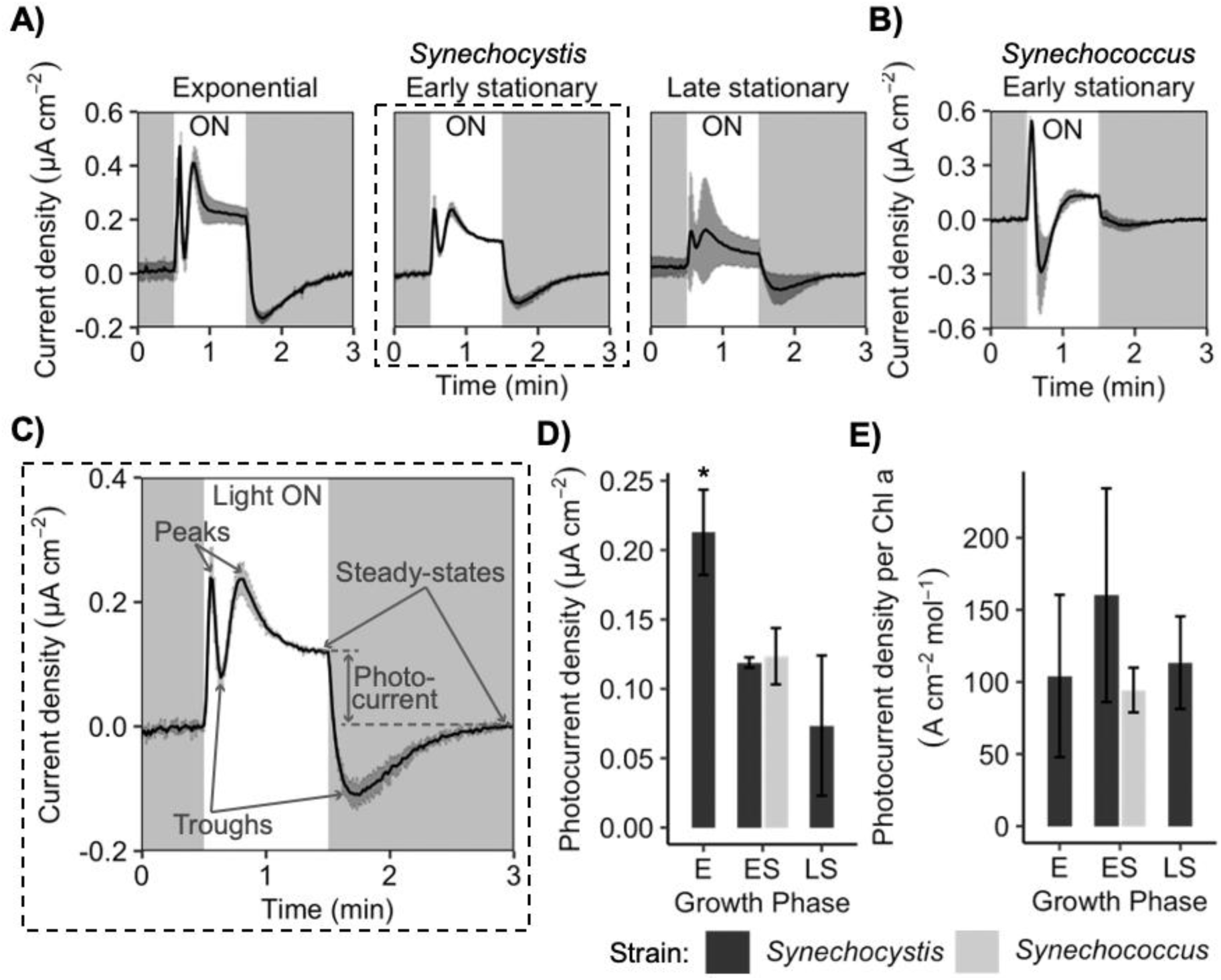
Photocurrent profiles of **A)** *Synechocystis* cells at different growth phases, **B)** *Synechococcus* cells at early stationary phase, and **C)** *Synechocystis* cells at early stationary phase (dashed box from panel A) with features labelled. ON = light on. Photocurrent outputs from *Synechocystis* cells at different growth phases and *Synechococcus* cells at early stationary phase **D)** calculated from photocurrent profiles in panels A and B, and **E)** normalised to chlorophyll (Chl *a*) content of cells remaining on the electrodes after photoelectrochemistry experiments. E = exponential, ES = early stationary, LS = late stationary. All chronoamperometry measurements were recorded at an applied potential of 0.3 V vs. SHE, and under atmospheric conditions at 25 °C in BG11 medium electrolyte. Light conditions used: λ = 680 nm, 50 µmol photons m^−2^ s^−1^ (approximately 1 mW cm^−2^ equivalent). Data presented as mean ± standard deviation of three biological replicates, * = P < 0.05 compared to *Synechocystis* cells at early stationary phase using a one-way ANOVA and Tukey HSD.

A range of cyanobacterial species have been used in studies of photocurrent output, including in studies of isolated photosynthetic proteins and cells, making it difficult to draw comparisons.^6^ To investigate exoelectrogenesis in different model cyanobacterial species, we compared the photoelectrochemistry of early stationary phase cultures of *Synechocystis* and *Synechococcus* under the above protocol. *Synechococcus* also gave a reproducible, complex photocurrent profile that displayed some of the same features as that of *Synechocystis*, although the trough recorded at 12 s after illumination was significantly larger in *Synechococcus* than *Synechocystis*, obscuring where there was a peak 25 s after illumination in the photocurrent profile of *Synechocystis* (**Fig. 2B**). *Synechococcus* yielded photocurrent outputs (raw and chlorophyll-normalised, **Fig 2 D, E**) similar to those of *Synechocystis*. The individual features of the photocurrent profile may be attributable to waves of contribution from different proteins, for example. Since we found that the magnitude of the features in the photocurrent profile varied between the two cyanobacterial species, subsequent comparisons of the photoelectrochemistry of subcellular fractions, in order to localise the molecular origin of these features, utilised material derived only from *Synechocystis* cells.

### Preparation of subcellular fractions on IO-ITO electrodes

To identify the role of cyanobacterial cell structure in exoelectrogenesis, specifically to determine which outer layers were responsible for the complexity in the photocurrent profile of whole wild-type cells, we took a top-down approach. We analysed the photoelectrochemistry of subcellular fractions of *Synechocystis* cells, from which different outer layers had been removed (**Supplementary Table 1**).

To investigate the role of extracellular appendages, several previously characterised *Synechocystis* mutants lacking different components of the type IV pilus machinery were tested (**Fig. 3A**).^35^ These include the Δ*pilA1* and Δ*pilA*9*-slr2019* mutants, which lack the major or minor pilins, respectively, and the Δ*pilB1* mutant, which lacks all pili due to loss of the pili extension motor. The Δ*pilT1* mutant was also tested, which is hyperpiliated (i.e. pili permanently extended) due to loss of the pili retraction motor. The genotypes of the pilus mutants were confirmed by PCR (**Supplementary Fig. 2**).

**Figure 3.**
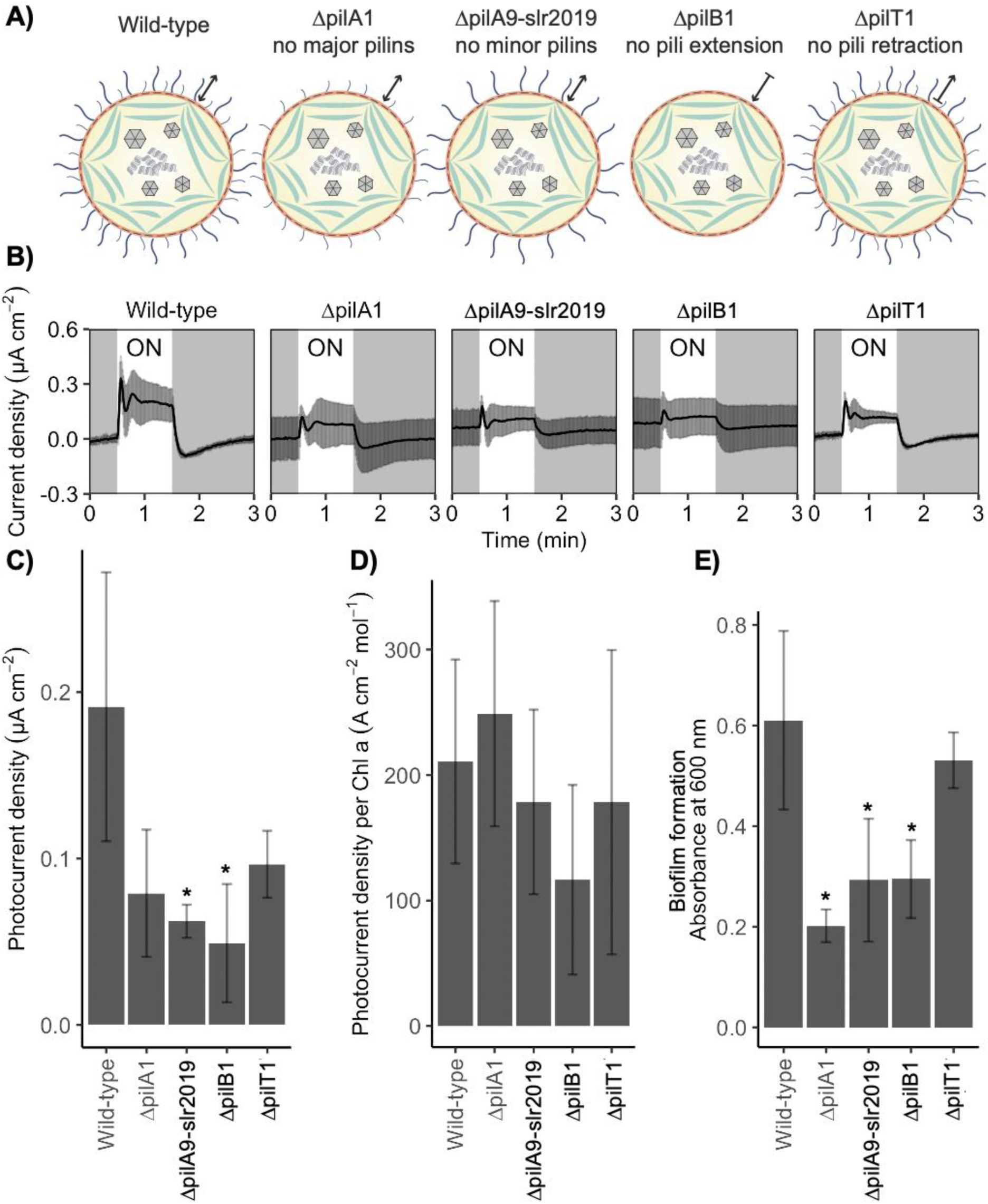
**A)** Schematic representation of *Synechocystis* pilus mutants with different components of the type IV pili machinery analysed in this study: wild-type cells, ΔpilA1 mutant cells lacking the major pili, ΔpilA9-slr2019 mutant cells lacking the minor pili, ΔpilB1 mutant cells lacking major and minor pili, and ΔpilT1 mutant cells that are hyperpiliated. Arrow indicates function of the pili extension/retraction mechanism. **B)** Photocurrent profiles of *Synechocystis* pilus mutants. ON = light on. Photocurrent outputs from *Synechocystis* pilus mutants **C)** calculated from photocurrent profiles in panel B, and **D)** normalised to chlorophyll (Chl *a*) content of cells remaining on the electrodes after photoelectrochemistry experiments. All chronoamperometry measurements were recorded at an applied potential of 0.3 V vs. SHE, and under atmospheric conditions at 25 °C in BG11 medium electrolyte. Light conditions used: λ = 680 nm, 50 µmol photons m^−2^ s^−1^ (approximately 1 mW cm^−2^ equivalent). **E)** Crystal violet assay of biofilm formation by *Synechocystis* pilus mutants loaded as per the protocol for photoelectrochemistry on ITO-PET. Data presented as mean ± standard deviation of three biological replicates for photoelectrochemistry and four biological replicates for biofilm assay, * = P < 0.05 compared to Wild-type using a one-way ANOVA and Tukey HSD.

To investigate the role of the most peripheral layer of the cell, the surface layer, a *Synechocystis* mutant lacking the surface layer (ΔS-layer) was tested (**Fig. 4A**). The ΔS-layer mutant was generated by using a recombination-based method to disrupt the *sll1951* gene that encodes the protein that solely constitutes the surface layer,^36^ and this was confirmed by PCR (Supplementary Fig. 3). Negative stain transmission electron microscopy of thin sections was used to confirm the absence of the surface layer in ΔS-layer cells (**Supplementary Fig. 4**). This showed the cell wall of the ΔS-layer mutant was thinner than that of the wild-type, and the most peripheral layer of the ΔS-layer mutant was smooth like lipopolysaccharide rather than the rougher glycosylated surface-layer present in wild-type cells. The appearance of the mutant was consistent with other mutants generated previously that lack the surface layer.^21^

**Figure 4.**
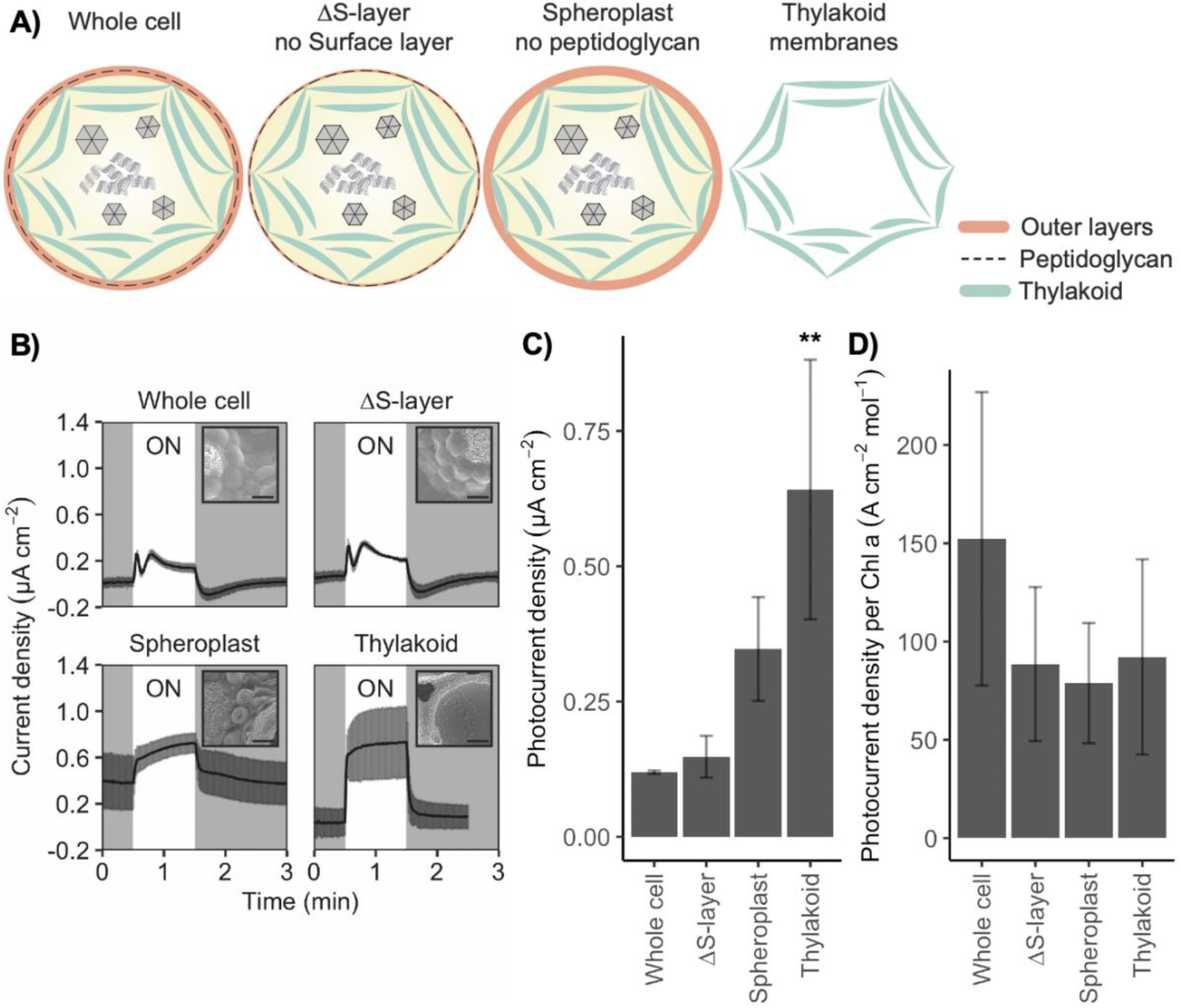
**A)** Schematic representation of subcellular fractions of *Synechocystis* cells analysed in this study: whole cells, mutant cells lacking the Surface layer (ΔS-layer); spheroplasts without peptidoglycan, and at least partial loss of the outer membrane and periplasmic space; and enriched thylakoid membranes. Pili are not depicted for clarity. **B)** Photocurrent profiles of subcellular fractions of *Synechocystis*. ON = light on. Inserts are scanning electron micrographs of each subcellular fraction loaded onto an IO-ITO electrode. Scale bar is 2 µm. Photocurrent outputs from subcellular fractions of *Synechocystis* **C)** calculated from photocurrent profiles in panel B, and **D)** normalised to chlorophyll (Chl *a*) content of cells remaining on the electrodes after photoelectrochemistry experiments. All chronoamperometry measurements were recorded at an applied potential of 0.3 V vs. SHE, and under atmospheric conditions at 25 °C in BG11 medium electrolyte for whole cells and ΔS-layer cells, spheroplast buffer for spheroplasts, and thylakoid buffer for thylakoid membranes. Light conditions used: λ = 680 nm, 50 µmol photons m^−2^ s^−1^ (approximately 1 mW cm^−2^ equivalent). Data presented as mean ± standard deviation of three biological replicates, ** = P < 0.01 compared to whole cells using a one-way ANOVA and Tukey HSD.

To investigate the role of the middle layers of the cell wall, spheroplasts, which do not have an intact peptidoglycan layer in the periplasmic space, were tested (**Fig. 4A**). Spheroplasts from Gram-negative bacteria, including cyanobacteria, also have a partial to complete loss of the outer membrane, which reduces the integrity of the periplasmic space in addition to the dissolution of the peptidoglycan layer.^37^ To generate spheroplasts from *Synechocystis*, whole wild-type cells from cultures that had reached early stationary phase were given a gentle lysozyme treatment.^38^ The preparation of osmotically fragile spheroplasts was confirmed by lysing the preparation in water and detecting phycobiliprotein in the supernatant by UV-Vis spectroscopy (**Supplementary Fig. 5**) and visualising the spheroplast ‘ghosts’ by light microscopy (**Supplementary Fig. 6**).

We also tested thylakoid membranes from *Synechocystis*, without any cytoplasm or bounding outer layers **(Fig. 4A)**. To isolate thylakoid membranes, whole wild-type cells from cultures that had reached early stationary phase were lysed and the membranes enriched by differential centrifugation of the lysate.^39^ Note that electrons would only be collected from the cytoplasmic face of the thylakoid membrane vesicles, so ‘inside-out’ thylakoid membrane vesicles should not contribute to or decrease the photocurrent output.

To confirm that the subcellular fractions had similar photosystem II activity, which generates the electrons from water oxidation that are eventually exported by the cell in exoelectrogenesis, we measured oxygen evolution. Whole wild-type cells, ΔS-layer mutant cells and spheroplasts showed similar photosystem II activity normalised to chlorophyll (whole wild-type cells: 91 ± 22 µmol(O_2_) mg^−1^(chl) h^−1^). Thylakoid membranes showed lower photosystem II activity (30 ± 6 µmol(O_2_) mg^−1^(chl) h^−1^) (**Supplementary Fig. 7**).

IO-ITO electrodes were loaded with the different subcellular fractions, and the incubation time was individually optimised to allow for penetration and adhesion of the material onto the electrodes. Loaded electrodes were rinsed in electrolyte optimal for each subcellular fraction (**Supplementary Table 2**) to remove excess biological material, then transferred to fresh electrolyte in the photoelectrochemical cell for chronoamperometry under light/dark cycles. The electrolyte for wild-type and mutant cells (ΔS-layer and pilus mutants) was BG11 medium, the electrolyte for spheroplasts was spheroplast buffer, and the electrolyte for thylakoid membranes was thylakoid buffer.

To confirm the integrity of the different subcellular fractions on the electrodes before photoelectrochemistry experiments, loaded electrodes were visualised by scanning electron microscopy (**Supplementary Fig. 8**). Wild-type and mutant cells (ΔS-layer and pilus mutants) as well as spheroplasts were observed as spheres on the electrodes and thylakoid membranes formed thin films on the surface of the electrodes. To visualise the integration and distribution of the subcellular fractions within the hierarchically structured IO-ITO electrodes, subcellular fractions were loaded on microscopy-dishes with an IO-ITO electrode on the coverslip bottom as per the protocol for photoelectrochemistry experiments. Chlorophyll autofluorescence of the subcellular fractions on the loaded electrodes was visualised by confocal microscopy (**Supplementary Fig. 9**) and confirmed that the subcellular fractions penetrated the IO-ITO electrodes through the channels and pores.

### Type IV pili are involved in cell-electrode adherence but do not contribute to exoelectrogenesis

To investigate whether components of the type IV pili have a role in exoelectrogenesis and the complexity of the photocurrent profile of cells, we measured the photocurrent profile of the different *Synechocystis* pilus mutants (**Fig. 3B**) under the standardised photoelectrochemical conditions used to capture the photocurrent profile of whole wild-type cells (**Fig. 2C**). All the pilus mutants exhibited complex photocurrent profiles similar to those of whole wild-type cells with two peaks at approximately 5 s and 25 s after illumination, a trough that reached its minimum approximately 12 s after illumination, and a broader trough that reached its minimum approximately 15 s after the light was turned off.

The photocurrent outputs of the different *Synechocystis* pilus mutants were calculated from the photocurrent profiles (**Fig. 3C**). The mutants lacking pili yielded significantly lower photocurrent output (e.g. Δ*pilB1* mutant cells approximately four-fold lower at 0.05 ± 0.03 µA cm^−2^) compared to their background wild-type cells (0.19 ± 0.08 µA cm^−2^). However, there was no statistically significant difference when the photocurrent outputs were normalised to the chlorophyll content remaining on the electrodes after photoelectrochemistry experiments (**Fig. 3D**).

To determine whether this was due to differences in cell loading, we measured the biofilm formation of wild-type and pilus mutant cells on flat ITO-coated polyethylene terephthalate (ITO-PET) using a crystal violet assay (**Fig. 3E**). The wild-type and hyperpiliated Δ*pilT1* mutant cells had significantly greater biofilm formation on ITO-PET (Abs_600_ = 0.6 ± 0.2 and 0.53 ± 0.06, respectively) than mutants lacking pili (e.g. Δ*pilA1* mutant cells approximately three-fold lower Abs_600_ = 0.20 ± 0.03). This confirms that the type IV pili are important in biofilm formation and surface adherence of *Synechocystis* onto materials, including ITO electrodes. Together with the photocurrent profiles, this indicates that in *Synechocystis* the type IV pili have no role in extracellular electron transfer from the cell to the electrode, but their loss results in lower photocurrent outputs because of a reduction in the number of cells adhering to the electrode.

### The periplasmic space determines the complex photocurrent profile of *Synechocystis* cells

Having shown that the type IV pili extracellular appendages were not responsible for the complexity in the photocurrent profile of cells, we considered the outer layers of the cyanobacterial structure. We measured the photocurrent profile under our standardised photoelectrochemical conditions of the subcellular fractions of *Synechocystis* with different outer layers removed (**Fig. 4B**). The ΔS-layer mutant (no surface layer) exhibited a complex photocurrent profile similar to that of whole wild-type cells. However, there was no difference in raw or chlorophyll-normalised photocurrent outputs between ΔS-layer mutant and the corresponding wild-type cells (**Fig. 4C, D**). Further, the ΔS-layer mutant showed similar amounts of biofilm formation to wild-type cells on ITO-PET (**Supplementary Fig. 10**). Therefore, we concluded that in *Synechocystis* the surface layer does not affect cell-electrode extracellular electron transfer or adherence.

The spheroplasts (compromised periplasmic space, from no peptidoglycan layer and incomplete outer membrane) and thylakoid membranes (no cell wall at all) showed monophasic photocurrent profiles under the same experimental regime, with a sharp increase in the current that reached a steady-state a few seconds after illumination, and a sharp decrease in the current that reached a steady-state a few seconds after the light was turned off (**Fig. 4B**). In addition, the steady-state current in the dark was higher in spheroplasts than whole cells. There was no significant difference in chlorophyll-normalised photocurrent densities between spheroplasts or thylakoid membranes and whole cells (**Fig. 4D**). We accounted for differences in photocurrent profile that would be caused by different electrolytes, pH, buffering and loading techniques (**Supplementary Fig. 11-13**). This indicates that the periplasmic space is responsible for the complex features of the photocurrent profile in cells.

### Comparison of wiring efficiency and longevity of subcellular fractions

To determine how directly wired the different subcellular fractions were to the electrodes, we measured the effect of adding an exogenous electron mediator that enables efficient indirect electron transfer from most active photosynthetic electron transport pathways present at the surface of the electrode (independent of the spatial arrangement, for example). We performed the chronoamperometry experiments in the presence of the model mediator 2,6-dichloro-1,4-benzoquinone (DCBQ) at an applied potential of 0.5 V vs SHE for maximal mediation by DCBQ (E*_m_* = 0.315 V vs SHE).^13^ We calculated the increase in photocurrent output in the presence of DCBQ relative to the photocurrent without the exogenous mediator added. If the subcellular fraction was poorly wired, then DCBQ would significantly increase the photocurrent output; whereas if the subcellular fraction was already wired well, then DCBQ would have a negligible effect on the photocurrent output. DCBQ gave rise to an approximately 60-fold higher photocurrent current for whole cells, whereas it gave rise to a three-fold higher photocurrent current for enriched thylakoid membranes (**Fig. 5A**). The fact that addition of the exogenous mediator resulted in a bigger increase in photocurrent output for whole cells than for thylakoid membranes indicates that enriched thylakoid membranes were inherently more efficiently wired to the electrodes than whole cells. The wiring of the spheroplasts was intermediate between whole cells and thylakoid membranes.

**Figure 5.**
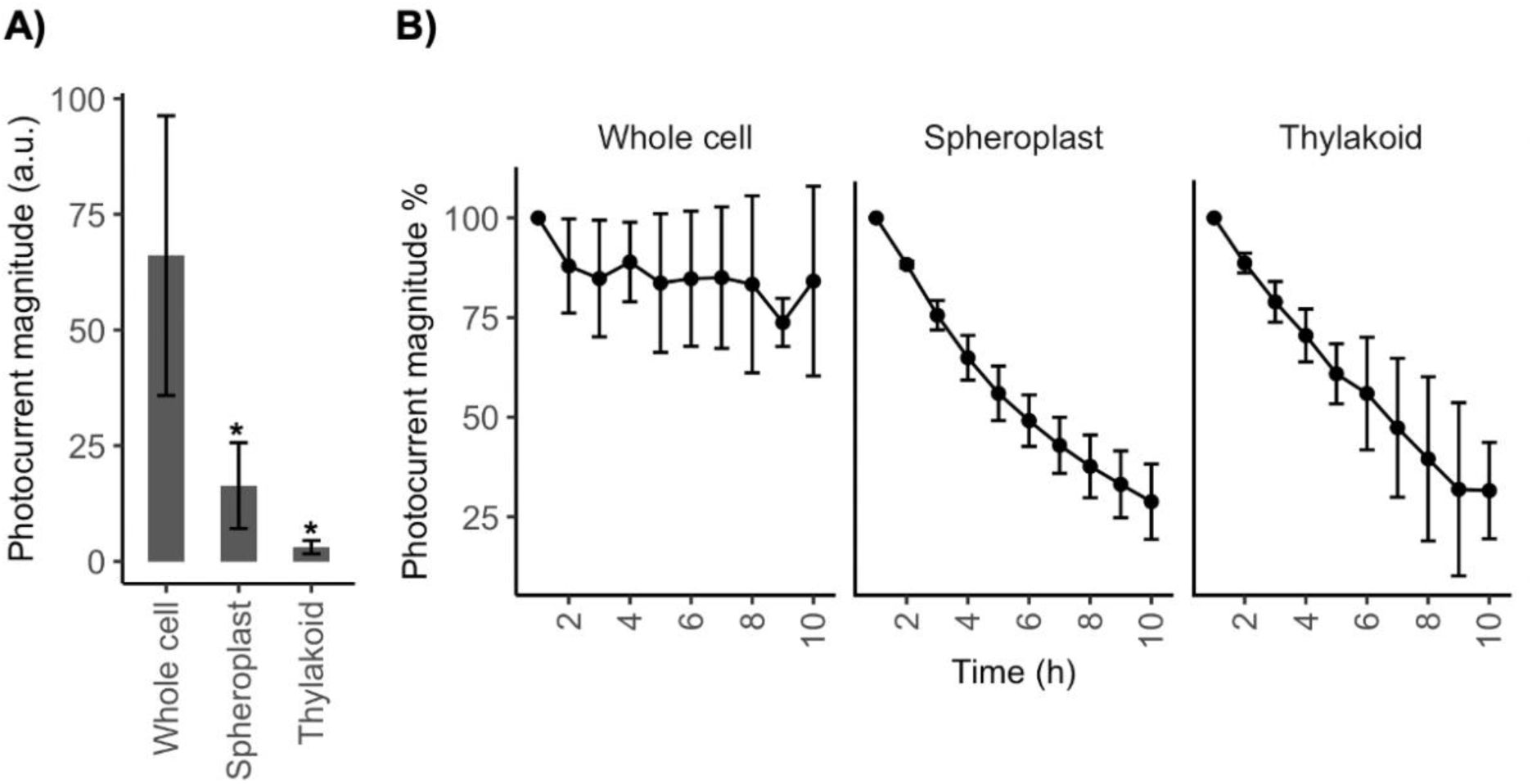
**A)** Photocurrent outputs from subcellular fractions of *Synechocystis* with DCBQ (1 mM) electron mediator relative to their photocurrent output with no exogenous mediator added as 1 arbitrary unit (a.u.). Chronoamperometry measurements with DCBQ were recorded at an applied potential of 0.5 V vs. SHE. Data presented as mean ± standard deviation of three biological replicates, * = P < 0.05 compared to whole cells using a one-way ANOVA and Tukey HSD. **B)** Photocurrent outputs from subcellular fractions of *Synechocystis* over 10 h as a percentage of the initial photocurrent output. Longevity chronoamperometry measurements were recorded at an applied potential of 0.3 V vs. SHE. Data presented as mean ± standard deviation of three biological replicates. All chronoamperometry measurements were recorded under atmospheric conditions at 25 °C in BG11 medium electrolyte for whole cells, spheroplast buffer for spheroplast and thylakoid buffer for thylakoid membranes. Light conditions used: λ = 680 nm, 50 µmol photons m^−2^ s^−1^ (approximately 1 mW cm^−2^ equivalent).

To determine the long-term stability of the different subcellular fractions, which is an important consideration in practical applications, we measured the photocurrent outputs of the different subcellular fractions over a 10 h period. We calculated the photocurrent output each hour relative to the initial photocurrent output (**Fig. 5B**). The photocurrent output of whole cells remained approximately the same across the 10 h period, consistent with previous studies.^13^ On the other hand, the photocurrent outputs of spheroplasts and thylakoid membranes fell to approximately 30% of their initial photocurrent outputs by the 10^th^ hour. Spheroplasts and thylakoid membranes had similar photocurrent output half-lives of approximately 6 h (spheroplasts 49 ± 6 %, thylakoid membrane 56 ± 14% initial photocurrent output at the 6^th^ hour). Thylakoids performed similarly to previous studies, likely due to protein degradation or photodamage that could not be repaired outside a living cell.^18^ Although one might have expected spheroplasts to perform for longer, their performance is consistent with the reported ability of spheroplasts of other bacteria to carry out metabolic reactions for only a few hours and be unable to replicate normally.^40, 41^

## Discussion

In this study, we were able to observe complex photocurrent profiles of cyanobacterial cells reproducibly under standardised microbiological and analytical photoelectrochemical conditions. While complex photocurrent profiles have been recorded in other studies,^8, 11–16^ drawing comparisons between them is challenging due to the use of different conditions. We kept constant, among other things, the ITO electrode material with an inverse-opal high surface area, the temperature, the intensity and wavelength of light (and used physiologically relevant light intensity), and the amount of chlorophyll loaded onto the electrodes. We also quantified the chlorophyll remaining on the electrodes after experiments, enabling comparison of photocurrent outputs normalised to chlorophyll. We then were able to systematically investigate the effect of growth phase, cyanobacterial species, and outer layers of the cell structure on exoelectrogenesis by varying those parameters in the biological samples we analysed using our robust photoelectrochemistry protocol. We found that the growth phase of the cells did not have a significant effect on exoelectrogenesis, however the photocurrent profiles of the two different model cyanobacterial species and some subcellular fractions differed.

While the photocurrent profiles of the two model organisms *Synechocystis* and *Synechococcus* exhibit complexity and the same broad features, we observed differences in the magnitudes of their features. For example, the trough recorded at 12 s after illumination was significantly deeper and broader in *Synechococcus* than *Synechocystis*, engulfing the peak at 25 s after illumination. A possible explanation is the different shapes of the cells, as *Synechocystis* and *Synechococcus* are spherical and rod-shaped, respectively, and cell morphology has been predicted to have a role in exoelectrogenesis.^42^ But the exact reason for the difference in their photocurrent profiles remains to be determined and will be the subject of future studies examining other model organisms.^43^

We observed that spheroplasts, without an integrous periplasmic space,^37^ yielded a monophasic photocurrent profile distinct from the complex profile of whole cells. This indicates that the periplasmic space contributes significantly to the complexity in the photocurrent profile of whole cells and likely acts as a gate for electron flow. The periplasmic space in *Synechocystis* contains redox active metabolites^44^ and proteins involved in reductive metal acquisition^45^, such as FutA2, which binds Fe^3+^ prior to uptake by the FutB/C plasma membrane transporter and is also proposed to be involved in copper-transport,^46, 47^ and CopM, which binds Cu^+^.^48^ There is evidence that reductive iron acquisition is linked to the respiratory terminal oxidases,^49^ and both iron acquisition and oxidase activity have been shown to be involved in exoelectrogenesis.^3, 4^ In addition, the periplasmic space bounded by the outer membrane provides a controlled environment on the extracellular side of the plasma membrane, which contains many proteins and a plastoquinol pool from the partial respiratory electron transfer chain.^45, 50, 51^ Future studies should examine the role of individual components of the periplasmic space, and periplasmic-dependent components of the plasma membrane, on exoelectrogenesis in *Synechocystis*.

We also observed that enriched thylakoid membranes yielded a monophasic photocurrent profile, similar to that of isolated photosystem II reported using similar conditions.^13^ Isolated photosystem II participates in direct electron transfer to IO-ITO electrodes,^52^ and it seems likely that the photosystem II in the enriched thylakoid membranes is functioning similarly. Future studies should use analytical photoelectrochemistry to study further the electron transfer from thylakoid membranes to electrodes to inform us about the photosynthetic electron transport chain, respiratory electron transport chain and other redox reactions in the thylakoid membranes.

Cyanobacterial exoelectrogenesis has long been hypothesised to involve a contribution from direct extracellular electron transfer via type IV pili.^6, 13, 51^ Previous atomic force microscopy (AFM) studies, including by Gorby et al., reported that *Synechocystis* type IV pili are electrically conductive.^24, 25^ The pili of the heterotrophic exoelectrogen *Geobacter sp*. have been shown to be electrically conductive, although these are much smaller in diameter than those in *Synechocystis*, which also lack the hyper-aromaticity that facilitates conductivity.^53–55^ Another recent study by Thirumurthy et al. showed that *Synechocystis* type IV pili are not required for extracellular electron transfer, as mutant cells lacking type IV pili due to deletion of the *pilD* pilin gene (and another mutation in a *pilA* pilin protein processing gene) yielded similar photocurrent output to wild-type cells on a carbon cloth electrode.^15^ Further, the authors reported that the *Synechocystis* type IV pili are not electrically conductive as measured by AFM, although the sample was not under low inorganic carbon conditions as described by Gorby et al..^24^ In this study, we observed that a variety of pilus mutant cells yielded complex photocurrent profiles and similar photocurrent outputs (normalised to chlorophyll content) compared to wild-type cells. Therefore, we concluded that type IV pili, including the major and minor pilins or the pili retraction and extension mechanisms, do not contribute to exoelectrogenesis in *Synechocystis*, confirming the study by Thirumurthy et al..^15^ In addition, in this study we also observed that the surface layer did not contribute to or impede exoelectrogenesis in *Synechocystis*.

It remains to be determined if cyanobacterial exoelectrogenesis involves a contribution from direct extracellular electron transfer via non-pilus or surface-layer cell features. Heterotrophic exoelectrogens such as *Geobacter* sp., *Shewanella oneidensis* and *Pseudomonas putida* are known to have redox metabolites and proteins in their extracellular polysaccharide matrices.^56, 57^ In this study we did not specifically investigate the extracellular polysaccharide matrix, but this will be a subject of future studies.

Nevertheless, we found that the cell surface features of *Synechocystis* affected the cell loading on the electrodes. *Synechocystis* is known to live as a planktonic species, as well as undergoing flocculation or biofilm formation.^23, 58^ Type IV pili have been shown to have a role in biofilm formation in heterotrophic bacteria such as *Pseudomonas aeruginosa*.^59, 60^ In cyanobacteria, type IV pili have been shown to have a role in aggregation and flocculation.^35, 61^ Type IV pili were shown to be not involved in biofilm formation in *Synechococcus*,^62^ but involved in *Synechocystis* biofilm formation,^58^ both on glass, which is nonphysiological and not an electrode material. In this study, we observed that mutant cells with reduced pili were less adherent than wild-type or hyperpiliated cells on ITO electrode materials. Previous attempts to increase BPV outputs have focused on optimising the electrode and biological materials separately, for example. Our results suggest that future attempts should explore optimising the cell-electrode interface, including increasing cell adherence to the electrode and therefore increasing loading and decreasing the fragility of electrodes, reducing loss of non-adherent cells when operating. The hyperpiliated Δ*pilT1* mutant is a promising candidate, as are other mutants genetically engineered to have greater biofilm formation.^63^

The wiring efficiency of whole cells to electrodes could also be improved. Previous benchmark power outputs in two-electrode biophotovoltaic devices and photocurrent density outputs in three-electrode photoelectrochemical set-ups have come from increasing wiring through exogenous mediators, such as quinones,^64, 65^ phenazines^66^ or potassium ferricyanide,^7, 67^ or redox polymers.^68, 69^ However such exogenous mediators give rise to a redox potential drop during the electron transfer process, and may be cytotoxic and unsuitable for large-scale applications.^66, 70^ We have shown here that by modifying the outer layers of the cell structure the wiring can be improved, as we found that spheroplasts had intermediate wiring between whole cells and thylakoid membranes. However, spheroplasts did not maintain the good long-term stability of whole cells.

## Conclusion

This study demonstrated how analytical photoelectrochemistry performed on systematic microbiological samples is an effective approach for dissecting the role of biological components in exoelectrogenesis. We identified that the periplasmic space significantly contributed to the complex photocurrent profile cells, but the surface layer or type IV pili did not. This supports the existence of an endogenous electron mediator that facilitates indirect extracellular electron transfer from *Synechocystis* cells to the electrode. This approach opens up many possibilities for dissecting the role of specific biological components in extracellular electron transfer events by exoelectrogenic cells.

## Methods

### Cell Culture and Growth Conditions

*Synechocystis sp.* PCC 6803 (*Synechocystis*) and *Synechococcus elongatus sp.* PCC7942 (*Synechococcus*) were cultured photoautotrophically under 50 µmol photons m^−2^ s^−1^ of continuous white light at 30°C in BG11 medium supplemented with 10 mM NaHCO_3_.^71^ Liquid cultures were bubbled with air and shaken at 120 rpm. 1.5% (w/v) agar was used in solid medium. Antibiotics were added at a final concentration of 25 µg ml^−1^ chloramphenicol, 50 µg ml^−1^ kanamycin and 50 µg ml^−1^ apramycin as appropriate. Culture growth was measured by attenuance at 750 nm (OD_750_). Culture chlorophyll concentration (nmol(chl *a*) ml^−1^) was calculated from absorbances at 680 nm and 750 nm: (A_680_-A_750_) × 10.814.^72^ All absorbance measurements were taken using a UV-1800 Spectrophotometer (Shimadzu).

### Pilus mutants

The *Synechocystis* pilus mutants and Wilde Lab wild-type strain^35^ were a gift from Prof. Annegret Wilde (University of Freiburg, Germany) and Prof. Conrad Mullineaux (Queen Mary University of London, UK). The mutant lacking the major pilin (Δ*pilA1*) had the *pilA1* (*sll1694*) gene interrupted with a chloramphenicol resistance cassette.^35^ The mutant lacking the minor pilins (Δ*pilA9-slr2019*) had the *pilA9-slr2019* operon interrupted with a kanamycin resistance cassette.^35^ The mutant lacking all pili (Δ*pilB1*) used had the *pilB1* gene (*slr0063*) interrupted with a kanamycin resistance cassette, and the hyperpiliated mutant (Δ*pilT1*) had the *pilT1* gene (*slr0161*) interrupted with an apramycin resistance cassette.^35^ The genotypes were confirmed by PCR amplification of the interrupted loci in the pilus mutants and the wild-type background using screening primers in Supplementary Table 3, followed by agarose gel electrophoresis of the PCR products.

### Generation of the surface-layer knock out

The surface layer gene (*sll1951*) was disrupted in the Howe Lab wild-type strain of *Synechocystis*^73^ to generate a marker-less surface-layer mutant (ΔS-layer) by a method previously described.^36^ The genome sequence of *Synechocystis*^74^ was consulted via Cyanobase (http://genome.kazusa.or.jp/cyanobase) for primer design, primers used are listed in Supplementary Table 5 and PCR was performed by standard procedures using Phusion high fidelity DNA polymerase (NEB). Gene deletion of *sll1951* was performed by amplifying an upstream 966bp fragment in the N-terminal region of *sll1951* using primers Sll1951leftfor and Sll1951leftrev and a 954bp downstream fragment in the N-terminal region of *sll1951* using primers Sll1951rightfor and Sll1951rightrev, followed by insertion of the respective fragments into the *Eco*R1/*Bam*H1 and *Bam*H1/Xba1 sites of pUC19 to generate pSll1951-1. The *Bam*H1 digested *npt1*/*sacRB* cassette from pUM24Cm^75^ was inserted into the *Bam*H1 site between the upstream and downstream fragments in pSll1951-1 to generate pSll1951-2. To generate marked mutants approximately 1 µg of pSll1951-1 plasmid was mixed with *Synechocystis* cells for 4 hours in liquid media, followed by incubation on BG11 agar plates for approximately 24 hours. An additional 3 ml of agar containing kanamycin was added to the surface of the plate followed by further incubation for approximately 1-2 weeks. Transformants were subcultured to allow segregation of mutant alleles. Segregation was confirmed by PCR using primers Slayer_screen_F and Slayer_screen_R, which flank the deleted region. Generation of unmarked mutants was carried out according to a method previously described.^76^ To remove the *npt1*/*sacRB* cassette, the mutant line was transformed with 1 µg of the markerless pSll1951-2 construct. Following incubation in BG-11 liquid media for 4 days and agar plates containing sucrose for a further 1-2 weeks, transformants were patched on kanamycin and sucrose plates. Sucrose resistant, kanamycin sensitive strains containing the unmarked deletion were confirmed by PCR amplification using primers Slayer_screen_F and Slayer_screen_R, followed by capillary electrophoresis of the PCR products using a QIAxcel Advanced System (QIAGEN).

### Transmission electron microscopy

The phenotype of the ΔS-layer mutant was confirmed by thin-section and negative stain transmission electron microscopy of newly generated ΔS- layer mutant and wild-type background.

### Preparation of spheroplasts

Spheroplasts were prepared from 50 ml of *Synechocystis* culture of the Howe Lab wild-type strain^73^ grown to early stationary phase (OD_750_ = 1.0) according to a previously reported method,^38^ with the following alterations: the cell-lysozyme mixture was not sonicated and only incubated for 1 h at 30°C, and the spheroplasts were harvested by centrifugation at 3000 g for 7 min at room temperature, which was based on an earlier description of the procedure.^77^ The formation of spheroplasts was confirmed by resuspending the spheroplast preparation in water and visualising by light microscopy the ghost-like appearance of lysed spheroplasts resulting from the hypo-osmotic shock.^21^ As a further test that spheroplasts were prepared, the presence of phycobilin protein was measured by UV-Vis spectroscopy in the supernatant of a spheroplast preparation resuspended in water.

### Thylakoid membrane enrichment

Cyanobacterial thylakoid membranes were enriched from 1 L of *Synechocystis* culture of the Howe Lab wild-type strain^73^ grown to early stationary phase (OD_750_ = 1.0) according to a previously reported method,^39^ with the exception that the 1mM benzamidine in Buffer A was replaced with a 1x concentration of cOmplete™ Protease Inhibitor Cocktail. The thylakoid membrane enrichment was resuspended to a final volume of 200 µl with a fine brush. The chlorophyll concentration was measured by methanol extraction and UV-Vis spectroscopy, using the extinction co-efficient of Chl *a* at 665.5 nm in methanol (70020 [mol chl a]^−1^ dm^3^ cm^−1^).^78^

### Oxygen evolution measurements

Oxygen evolution was measured using a Clark Electrode consisting of an Oxygraph Plus Electrode Control Unit, S1 Oxygen Electrode Disc, DW2/2 Electrode Chamber and a LED1 High Intensity LED Light Source (Hansatech Instruments). Measurements were performed on 1.5 ml samples of each subcellular fraction of *Synechocystis* containing 10 µg ml^−1^ of Chl *a* in appropriate electrolyte supplemented with 1 mM 2,6-dichloro-1,4-benzoquinone (DCBQ) and 1 mM potassium ferricyanide.

Measurements were collected at 25°C with 1 min darkness, followed by 1 min of 1500 µmol photons m^−2^ s^−1^ light at 627 nm. The rate of oxygen production in the dark was subtracted from that in the light, and normalised to Chl *a* content. Data were collected from three biological replicates, each with three technical replicates.

### Photoelectrochemistry

Inverse opal ITO (IO-ITO) electrodes with 10 μm macropores and 3 μm interconnecting channels at a thickness of 40 µm were prepared using the method previously reported.^13^

For wild-type *Synechocystis* and *Synechococcus*, pilus mutants and the ΔS-layer mutant, planktonic cultures of early stationary phase cells at OD_750_ of ca. 1 (unless growth phase stated otherwise) were concentrated by centrifugation at 5000 g for 10 min, the supernatant removed and the pellet resuspended in fresh BG11 medium to a concentration of 150 nmol Chl *a* ml^−1^. 250 μL of this solution was dropcast onto the IO-ITO electrodes and left overnight at room temperature in a humid chamber in the dark to allow cell penetration and adhesion, yielding cell-loaded electrodes that were used for analysis 16 h later.

For spheroplasts, the final preparation was resuspended in spheroplast buffer to a concentration of 1500 nmol Chl *a* ml^−1^. 25 μL of this solution was dropcast onto the IO-ITO electrodes and left for 1 h at room temperature in a humid chamber in the dark to allow spheroplast penetration and adhesion, yielding spheroplast-loaded electrodes which were used immediately for analysis.

For isolated thylakoid membranes, the final preparation was resuspended in thylakoid buffer to a concentration of 225 nmol Chl *a* ml^−1^. 10 μL of this solution was dropcast onto the IO-ITO electrodes and left for 15 min at room temperature in a humid chamber in the dark to allow thylakoid membrane penetration and adhesion, yielding thylakoid-loaded electrodes which were used immediately for analysis.

All photoelectrochemical measurements were performed at 25°C using an Ivium Technologies CompactStat, with an Ag/AgCl (saturated) reference electrode (corrected by + 0.197 V for SHE), a platinum mesh counter electrode and loaded IO-ITO working electrode. Chronoamperometry experiments without an exogenous mediator added were performed at an applied potential 0.3 V vs SHE. Chronoamperometry experiments with 1 mM DCBQ exogenous mediator added were performed at an applied potential 0.5 V vs SHE.

Unless otherwise stated, photoelectrochemical measurements were performed in 4.5 ml electrolyte appropriate for the subcellular fraction. The electrolyte for isolated thylakoids was 50 mM MES buffer with, 15 mM NaCl, 5 mM MgCl, 2 mM CaCl_2_ (pH 6.0); the electrolyte for spheroplasts was 10 mM HEPES buffer with 10 mM MgCl_2_, 5 mM sodium phosphate and 0.5 M sorbitol (pH 7.5); the electrolyte for whole wild-type cells and mutant cells (ΔSlayer and pilus mutants) was BG11 medium (pH 8.5).

Chronoamperometry experiments were performed at a sampling rate 1 s^−1^, with light/dark cycles using a collimated LED light source (50 µmol photons m^−2^ s^−1^, approximately 1 mW cm^−2^ equivalent, 680 nm, Thorlabs). Photocurrent profiles were measured under 60 s light/90 s dark cycles, and when steady-state was reached, the following three photocurrent profiles were taken for analysis. Longevity experiments were performed under 59 min light/1 min dark cycles.

Photocurrents were normalised to the geometric area of the electrode (0.75 cm^2^) to obtain photocurrent densities. The Chl *a* content of the bio-loaded electrode was determined by scraping off the annealed ITO nanoparticles from the FTO coated glass into methanol (500 µL). The suspension was sonicated for 1 h in iced water, then centrifuged at 10 000 g for 3 min. The supernatant was analysed by UV-Vis spectroscopy and the Chl *a* concentration was determined using the extinction co-efficient of Chl *a* at 665.5 nm in methanol (70020 [mol chl a]^−1^ dm^3^ cm^−1^).^78^

For ‘wiring’ experiments, the photocurrent enhancement with 1 mM DCBQ exogenous mediator added was calculated relative to the photocurrent without an exogenous mediator added. For longevity experiments, the photocurrent at each hour was calculated as a percentage relative to the initial photocurrent.

Three biological replicates of each subcellular fraction were measured. The mean and standard deviation across three biological replicates was calculated. Photocurrent outputs were compared to early stationary phase cells, whole cell or wild-type as appropriate using a one-way one-way analysis of variance (ANOVA) and Tukey’s (honestly significant difference) (HSD) test.

### Scanning electron microscopy

Subcellular fractions were loaded onto IO-ITO electrodes and left for a set amount of time under darkness as per the protocol for photoelectrochemistry. The loaded electrodes were rinsed in the electrolyte appropriate for each subcellular fraction. The loaded electrodes were then fixed in liquid nitrogen-cooled ethane and freeze-dried overnight. The loaded electrodes were mounted on aluminium stubs with silver dark and coated with 15 nm iridium using an EMITECH K575X Peltier cooler. The sample was stored in a desiccator until imaged using a TESCAN MIRA3 FEG-SEM with a 30 kV beam acceleration. Images were processed using Illustrator.

### Confocal fluorescence microscopy

Subcellular fractions were loaded onto bespoke µ- dishes with an IO-ITO electrode on the coverslip bottom and left for a set amount of time under darkness as per the protocol for photoelectrochemistry. The loaded µ-dishes were washed once and then filled with the electrolyte appropriate for the subcellular fraction. The loaded µ-dishes were visualised using a Nikon Eclipse Ti2 Inverted Microscope. Brightfield and autofluorescence of chlorophyll images were taken. Images were processed using NIS Elements Viewer.

### Biofilm formation assay

A crystal violet assay was used to measure biofilm formation as adapted from a protocol previously described for *Synechocystis*.^58^ Pilus mutants and wild-type cells of the background used to construct the mutants (the Wilde laboratory strain) were loaded onto ITO-coated polyethylene terephthalate (ITO-PET) 1 cm^2^ discs in wells of a 24- well plate and left for under darkness as per the protocol for photoelectrochemistry. 150 µl was aspirated from each well without disturbing the biofilm before gently washing the cell-loaded disc thrice with 750 µl BG11 medium to remove non-adherent cells. Crystal violet solution (250 µl, 0.1% w/v) in dH_2_O was pipetted into each well and incubated for 15 min at room temperature. Following this, 200 µL was aspirated from each well and the disc was washed three times with 250 µl dH_2_O. 1 ml DMSO was pipetted into each well and the 24- well plate was vigorously shaken for 15 min. The absorbance at 600 nm of the crystal violet in the DMSO supernatant was measured and used as a proxy for the amount of cellular material bound to the ITO-PET.^79^ The surface layer mutant and the wild-type cells of the background used to construct the mutant (the Howe laboratory strain) were loaded onto ITO-PET in previously described biophotovoltaic devices^80^ and left under 50 µmol photons m^−2^ s^−1^ of continuous white light for five days before performing the crystal violet assay as above.

## Acknowledgements

The authors thank the Cambridge Trust (L.T.W.), the Biotechnology and Biological Sciences Research Council (BB/M011194/1 [J.M.L.]; David Phillips Fellowship BB/R011923/1 [to J.Z.Z., also supporting X.C.]) and the Waste Environmental Education Research Trust (D.J.L-S.) for financial support. The authors gratefully acknowledge Dr Karin Muller and Dr Jeremy Skepper of the Cambridge Advanced Imaging Centre for their assistance in this work in confirming the phenotype of the ΔS-layer mutant by Transmission Electron Microscopy, and Prof. Annegret Wilde (University of Freiburg, Germany) and Prof. Conrad Mullineaux (Queen Mary University of London, UK) for the gift of the pilus mutants used in this study.

## Author contributions

L.T.W. cultured cyanobacterial cells, prepared spheroplasts and performed photoelectrochemistry experiments on subcellular fractions. J.M.L. prepared thylakoids and performed photoelectrochemistry experiments on thylakoids. X.C. prepared the electrodes and performed the scanning electron microscopy experiments. L.T.W. performed confocal microscopy experiments. R.C. performed the biofilm formation assays. D.L.-S. generated the surface layer mutant. L.T.W. analysed the data. L.T.W., J.Z.Z. and C.J.H. wrote the manuscript. All authors contributed to the editing of this manuscript.

## Competing interests

The authors declare no competing interests.

## Additional information

The supplementary information document contains supplementary material, including supplementary figures and tables.

## Abbreviations

AFM: Atomic force microscopy
ANOVA: analysis of variance
BPV: biophotovoltaic device
Chl *a*: chlorophyll *a*
DCBQ: 2,6-dichloro-1,4-benzoquinone
E*_m_*: mid-point potential
ET: electron transfer
ETC: electron transport chain
Tukey HSD: Tukey’s honestly significant difference test
IO-ITO: inverse-opal indium-tin oxide
ITO-PET: indium-tin oxide coated polyethylene terephthalate
OD: optical density
S-layer: Surface layer
Synechococcus: Synechococcus elongatus PCC7942
*Synechocystis*: *Synechocystis sp*. PCC 6803
SHE: Standard hydrogen electrode

## Notes

### Competing Interest Statement

The authors have declared no competing interest.

