## Supplementary Information for "A biophotoelectrochemical approach to unravelling the role of cyanobacterial cell structures in exoelectrogenesis"

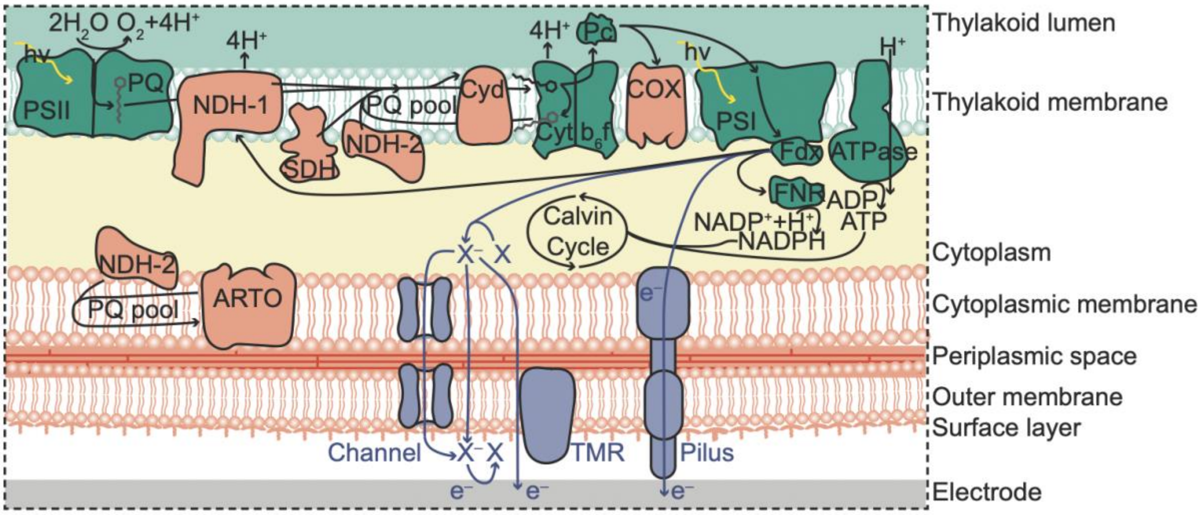


**Supplementary Figure 1** Schematic representation of the cell-electrode interface with arrows indicating the electron transfer (ET) pathways: known intracellular (black arrows)[1] and postulated ET pathways behind exoelectrogenesis (purple arrows).[2] 'X' is an unidentified electron carrier produced and exported by the cyanobacterial cells under illumination. Postulated mechanisms include electrons stemming from downstream of photosystem I (PSI) and passing to the outside of the cell via channels, passive diffusion, a putative transmembrane reductase (TMR), or type IV pili. Linear photosynthetic electron transport chain proteins (coloured green) are intermingled with the proteins (coloured orange) of the cyclic and respiratory electron transport chains in the thylakoid membrane, and there is a partial respiratory chain in the cytoplasmic membrane. The cell is bounded by the cell wall consisting of the cytoplasmic membrane, outer membrane, and surface layer. Type IV pili traverse these layers of the cell wall. The cytoplasmic and outer membranes are separated by a periplasmic space including the peptidoglycan layer. Proteins: Photosystem II (PSII), NAD(P)H dehydrogenase-like complexes 1 and 2 (NDH-1, -2), cytochrome $b_6f$ complex (Cyt $b_6$f), plastocyanin (Pc), photosystem I (PSI), ferredoxin (Fdx), ferredoxin-NADP$^+$ reductase (FNR), ATP synthase (ATPase), cytochrome *bd*-quinol oxidase (Cyd), cytochrome-*c* oxidase complex (COX), alternative respiratory terminal oxidase (ARTO). Membrane-bound mediators: plastoquinone/ plastoquinol (PQ/$PQH_2$). Phycobilisomes and flavodiiron proteins are not shown for clarity.

**Supplementary Table 1** The sub-cellular fractions of *Synechocystis* prepared for photoelectrochemistry analysis in this study.

| Whole wild-type cells starting material | Sub-cellular fraction | Description | Method of preparation | Ref. |
|---|---|---|---|---|
| Howe Group strain[3] | ∆S-layer mutant cells | Cells lacking the surface layer | Genetic removal of part of the *sll1951* gene that encodes the surface-layer protein | 4 |
| | Spheroplasts | Cells lacking an intact peptidoglycan layer in the periplasmic space | Gentle lysozyme treatment of whole wild-type cells | 5 |
| | Thylakoid membranes | Isolated thylakoid membranes | Thylakoid membranes enriched by differential centrifugation of wild-type cell lysate | 6 |
| Wilde Group strain[7] | ∆pilA1 mutant cells | Cells lacking major pili | Genetic removal of the *pilA1* (*sll1694*) gene that encodes the major pilin structural protein | 7 |
| | ∆pilA9-slr2019 mutant cells | Cells lacking minor pili | Genetic removal of the *pilA9-slr2019* operon that encodes the minor pilin structural proteins | 7 |
| | ∆pilB1 mutant cells | Cells lacking major and minor pili | Genetic removal of the *pilB1* (*slr0063*) gene that encodes the pili extension motor | 7 |
| | ∆pilT1 mutant cells | Hyperpiliated cells | Genetic removal of the *pilT1* (*slr0161*) gene that encodes the pili retraction motor | 7 |

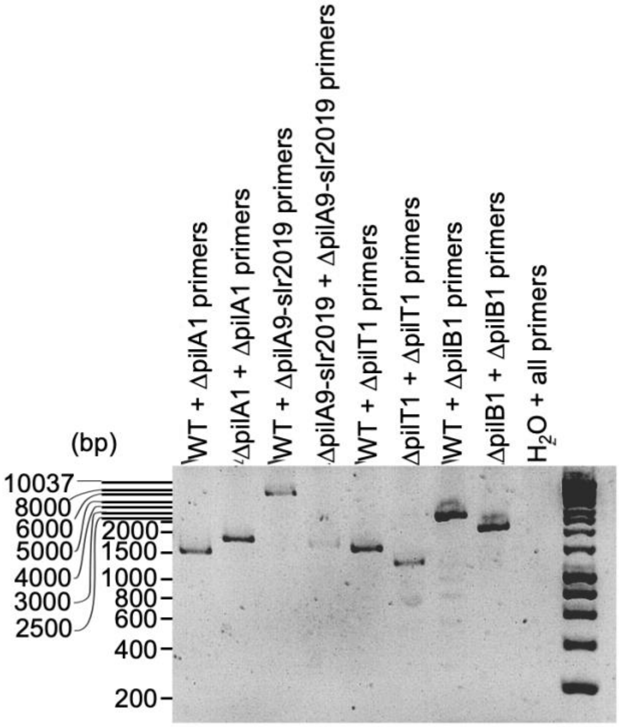


**Supplementary Figure 2** Agarose gel from electrophoresis of PCR products from amplification of the pilus gene locus in wild type and pilus mutant strains with appropriate primers (Supplementary Table 3). A negative control of autoclaved $dH_2O$ with all primers was used to demonstrate the lack of contamination. Far right lane is Bioline HyperLadder™ 1kb. All bands are at expected size (Supplementary Table 4).

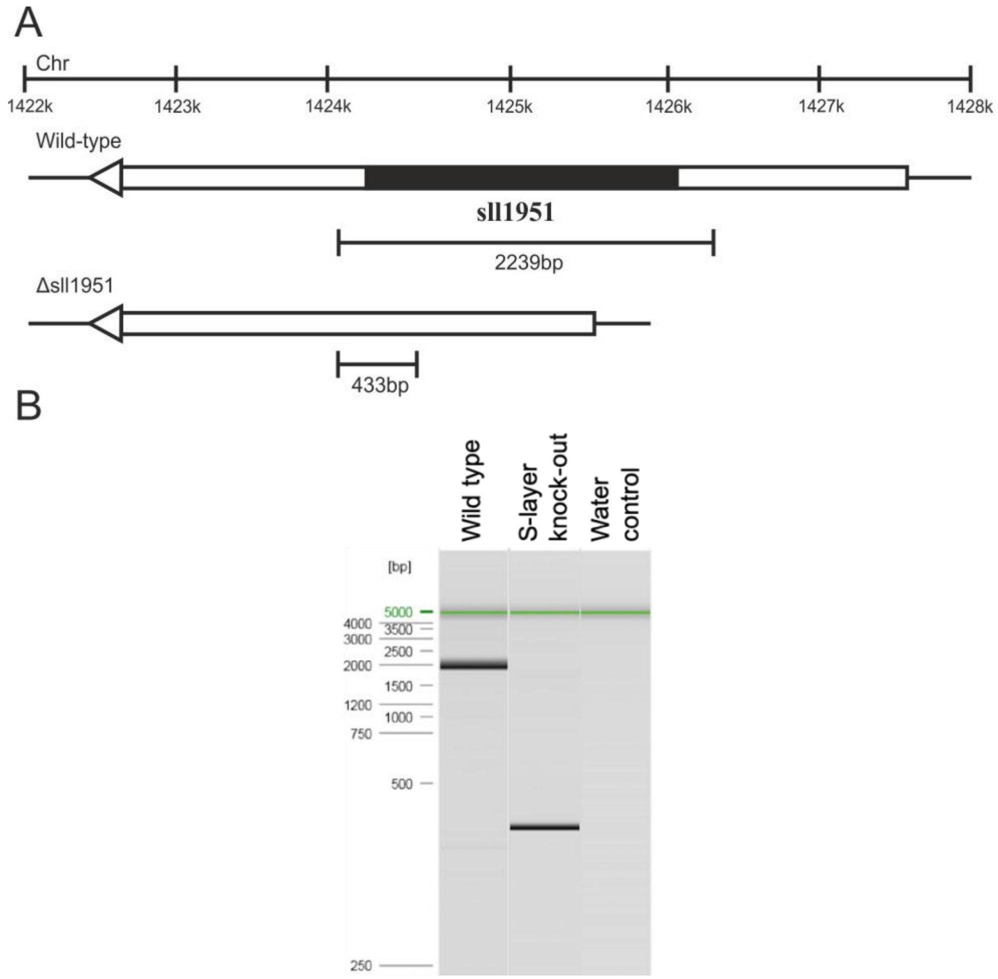


**Supplementary Figure 3 A)** Schematic representation of locus location in the *Synechocystis* genome (top) and the wild-type (middle) and unmarked surface layer knockout (bottom) profiles expected following amplification with primers flanking the deleted sequence. The region deleted in the mutant strain is shaded in black. **B)** Simulated gel from QIAxcel capillary electrophoresis of PCR products from amplification of the surface-layer gene locus in wild type and surface layer knock-out strains by primers Slayer_screen_F and Slayer_screen_R (Supplementary Table 5). The size of band expected for the wild type cells was 2239 bp and for the knock out 433 bp.

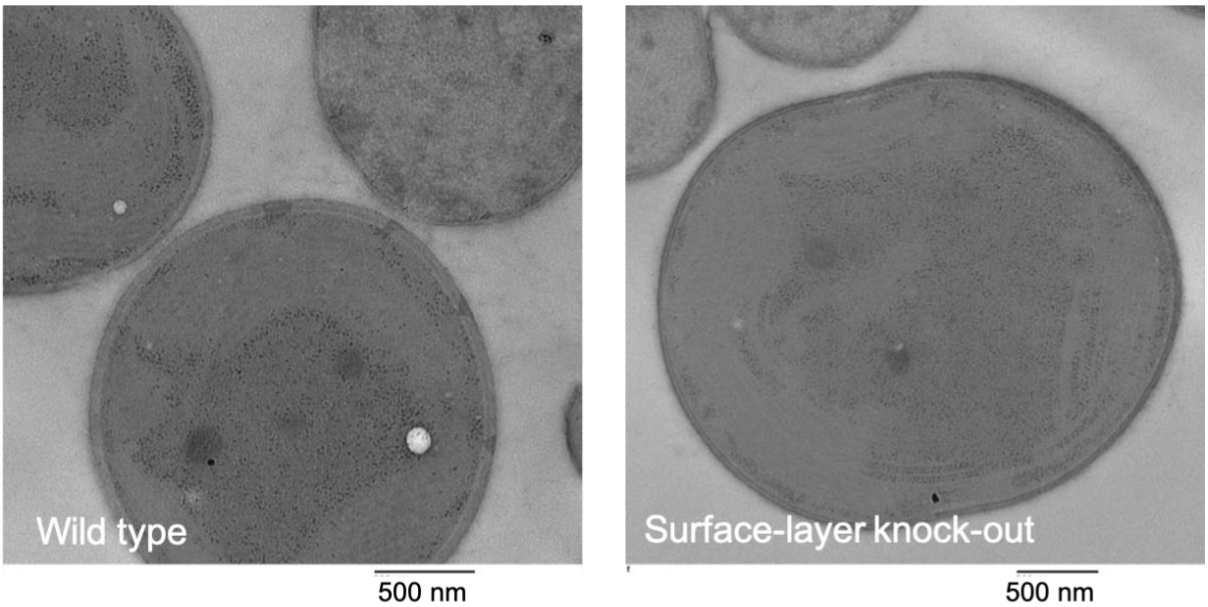


**Supplementary Figure 4** Transmission electron micrograph of wild type cells and surface-layer knock out cells. Scale bar 500 nm.

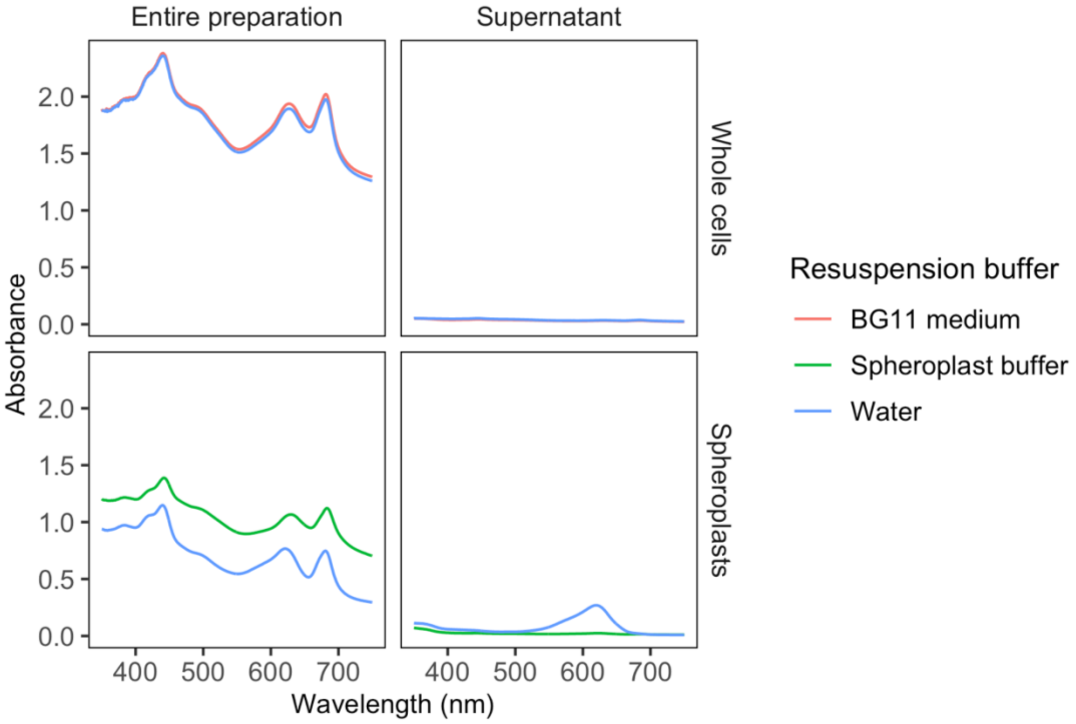


**Supplementary Figure 5** UV-Vis spectra of spheroplast preparations from 350 to 750 nm. Rows: whole cell and spheroplast preparations. Columns: entire preparation or the supernatant after centrifugation at 3000 g for 7 min. Samples were resuspended in either buffer appropriate for that fraction (BG11 medium for whole cells, or spheroplast buffer for spheroplasts) or water. Phycocyanin absorbs at a $\lambda_{max}$ = 622 nm.

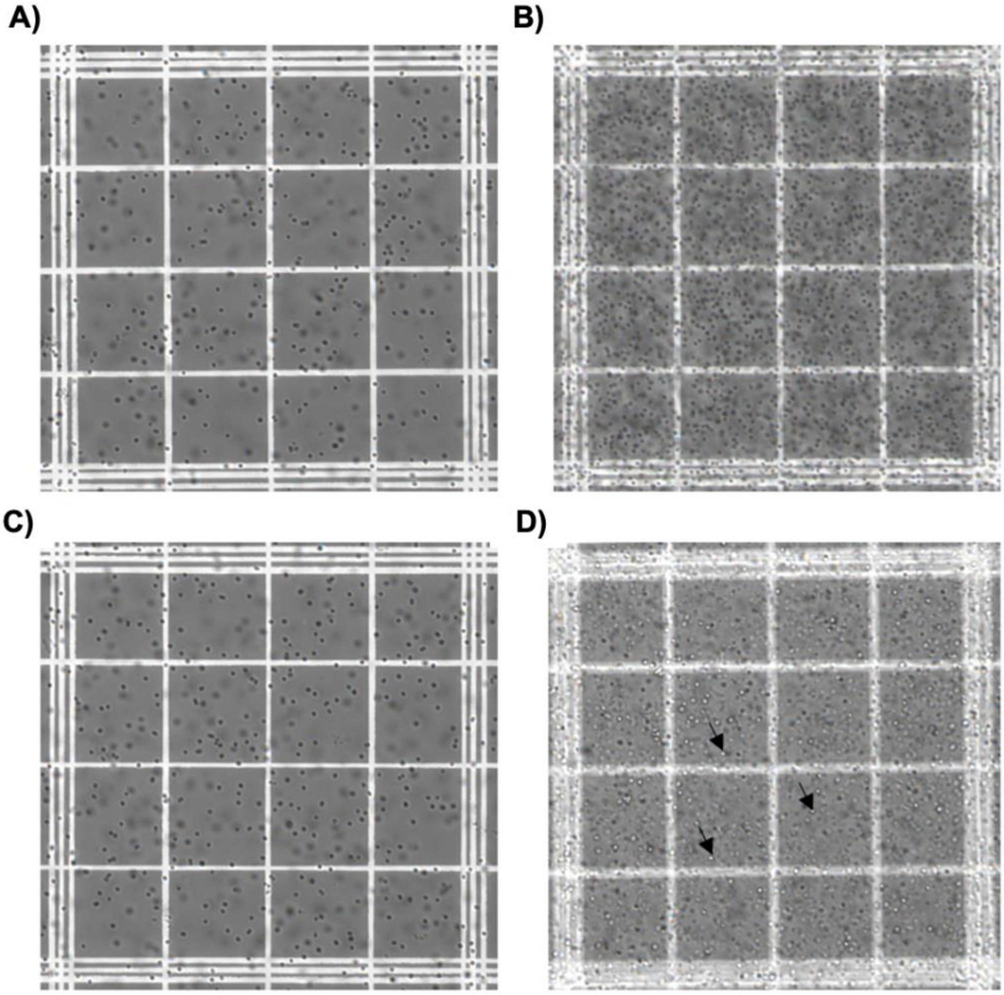


**Supplementary Figure 6** Light microscope images of spheroplast preparations at 20 x magnification of (A, C) whole cell 1/100 x dilution, and (B, D) spheroplast 1x preparations in (A, B) appropriate buffer (BG11 and spheroplast buffer, respectively) and (C, D) water. Some spheroplast ‘ghosts’ are indicated with arrows.

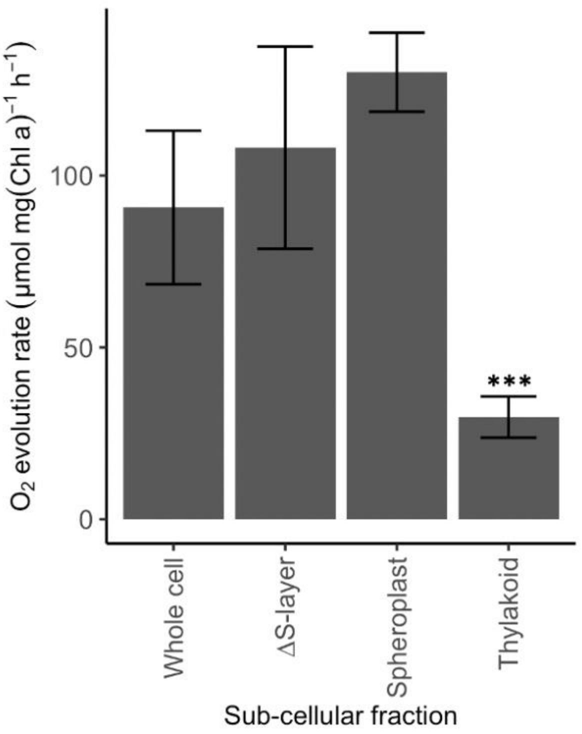


**Supplementary Figure 7** Oxygen evolution rates of different sub cellular fractions from Clark Electrode measurements using 1521 µmol photons $m^{-2}$ $s^{-1}$ light at 627 nm, 1 mM DCBQ and 1 mM potassium ferricyanide. The electrolyte for whole wild-type cells and surface-layer mutant cells (ΔS-layer) was BG11 medium (pH 8.5); the electrolyte for spheroplasts was 10 mM HEPES buffer with 10 mM $MgCl_2$, 5 mM sodium phosphate and 0.5 M sorbitol (pH 7.5); and the electrolyte for isolated thylakoids was 50 mM MES buffer with, 15 mM NaCl, 5 mM $MgCl_2$, 2 mM CaCl2 (pH 6.0). Data presented as the mean of three biological replicates, error bars are standard error in the mean, *** = $P < 0.001$ compared to whole cell using a one way ANOVA and Tukey post-hoc test.

**Supplementary Table 2** Key properties of buffers of each subcellular fraction

| **Subcellular fraction** | **Buffer** | **Osmolarity** | **pH** | **Buffering** | **Redox active species** |
|---|---|---|---|---|---|
| Wild-type cells and mutant cells (∆S-layer and pilus mutants) | BG11 medium | 58 mosmol | 8 | 175.1 µM phosphate buffer | $Mg^{2+}$, citric acid, $Mn^{2+}$, $Zn^{2+}$, $MoO_4^{2+}$, $Cu^{2+}$, $Co^{2+}$, ferric citrate |
| Spheroplast | Spheroplast buffer | 575 mosmol | 7.5 | 10 mM HEPES | $Mg^{2+}$ |
| Thylakoid membranes | Thylakoid buffer | 121 mosmol | 6.0 | 50 mM MES | $Mg^{2+}$ |

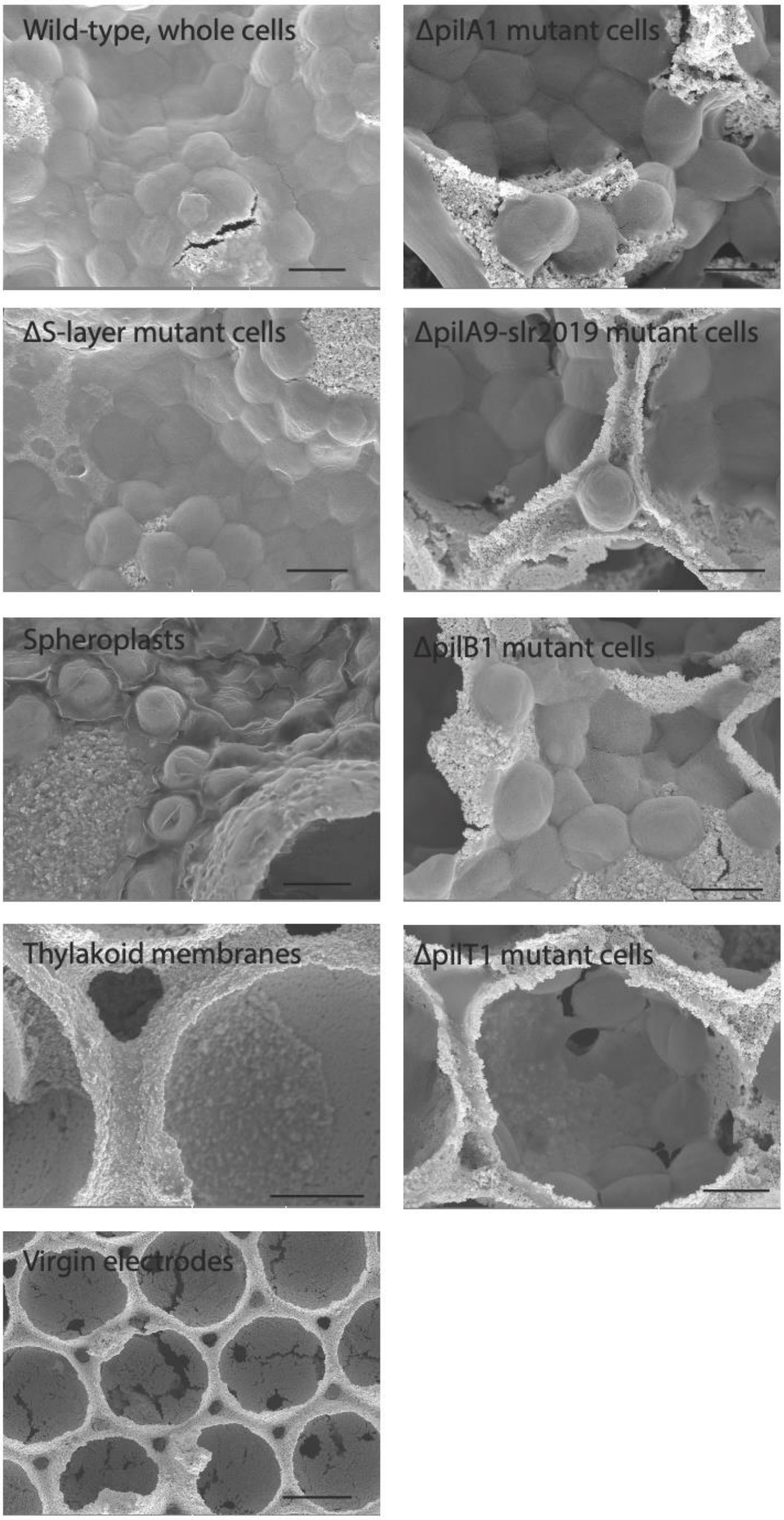


**Supplementary Figure 8** Scanning electron microscope images of sub-cellular fractions of *Synechocystis* loaded on IO-ITO electrodes according to the protocol for photoelectrochemistry. Scale bar is 2 µm, apart from the virgin electrodes where the scale bar is 5 µm.

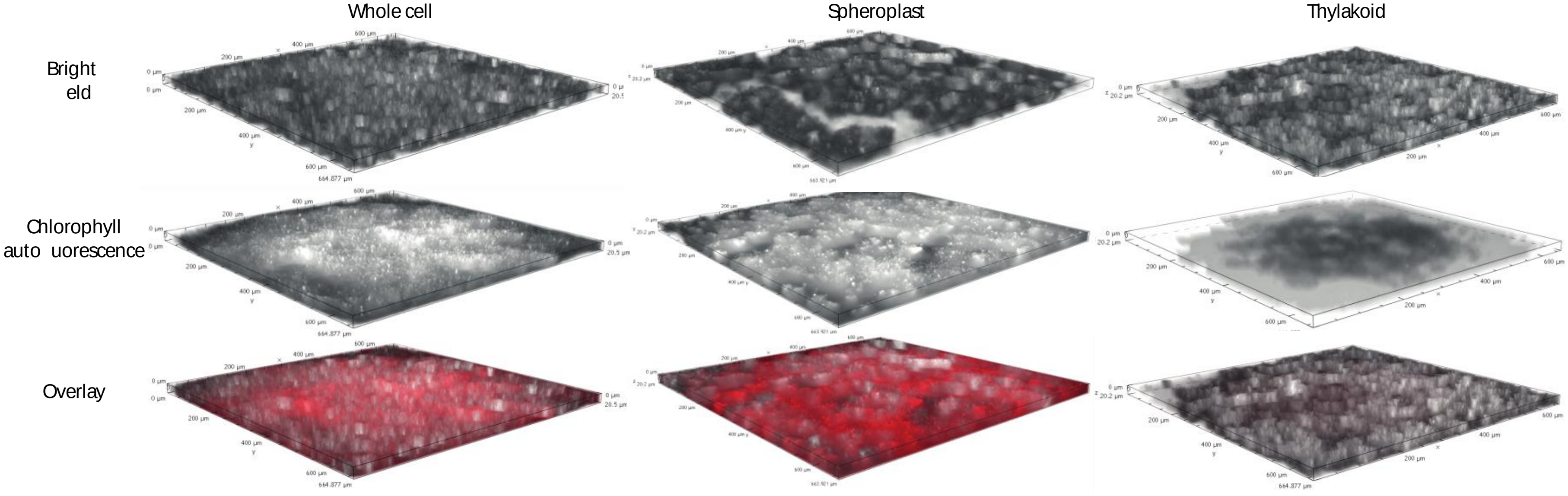


**Supplementary Figure 9** 3D visualisation of the sub-cellular fractions of *Synechocystis* integration and distribution within IO−ITO electrodes. The 3D view was reconstructed from Z-stacking images that were acquired by scanning 20 μm upward from the electrode base. Excitation: λex = 635 nm. Emission: λem = 690 nm.

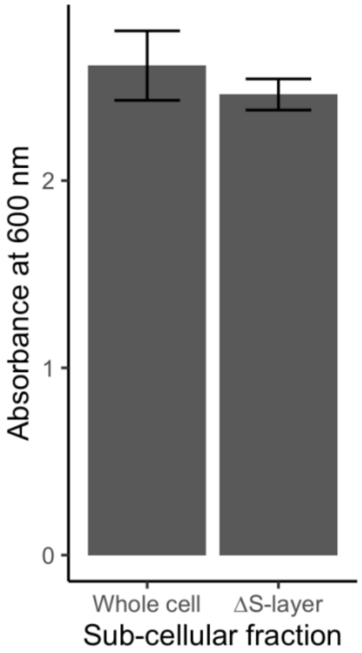


**Supplementary Figure 10** Crystal violet assay of *Synechocystis* wild-type, whole cells and surface layer mutant cells (∆S-layer). All loaded on ITO-PET. Data presented as the mean of four biological replicates, error bars are standard deviation, with no statistically significant difference using a one way ANOVA and Tukey HSD.

To confirm the difference in whole cell and spheroplast photocurrent profiles was not due to the different electrode loading techniques, whole cells were loaded on an electrode as per the spheroplast protocol and the photocurrent profile was measured. Whole cells still yielded a complex photocurrent profile under these conditions (**Supplementary Fig. 11**).

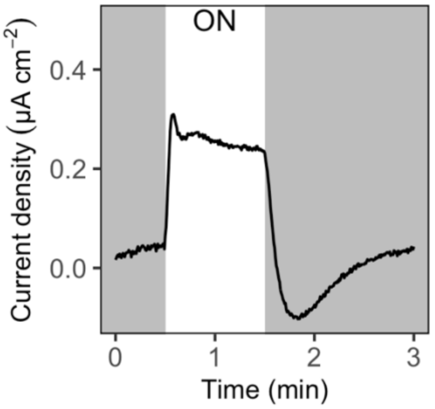


**Supplementary Figure 11** Photocurrent profile of whole wild-type *Synechocystis* cells loaded for 1 h as for spheroplasts. Chronoamperometry measurements were recorded at an applied potential of 0.3 V vs. SHE, and under atmospheric conditions at 25 °C in BG11 medium electrolyte. Light conditions used: λ = 680 nm, 50 µmol photons $m^{-2}$ $s^{-1}$ (approximately 1 mW $cm^{-2}$ equivalent). ON, light on. Data show one biological sample, presented as the mean of three consecutive light/dark cycles.

To confirm the difference in whole cell and spheroplast photocurrent profiles was not due to the different electrolytes (**Supplementary Table 2**), the photocurrent profile of whole cells was recorded in electrolytes with properties intermediate between BG11 medium and spheroplast buffer. Whole cells still yielded a complex photocurrent profile (**Supplementary Fig. 12**), although the kinetics of the photocurrent profile were different in the electrolytes at different pH. Incubation of whole cells in electrolyte identical to that used for spheroplasts or *vice versa* led to complete loss of exoelectrogenic activity, necessitating the use of buffers with intermediate properties (**Supplementary Fig. 13**).

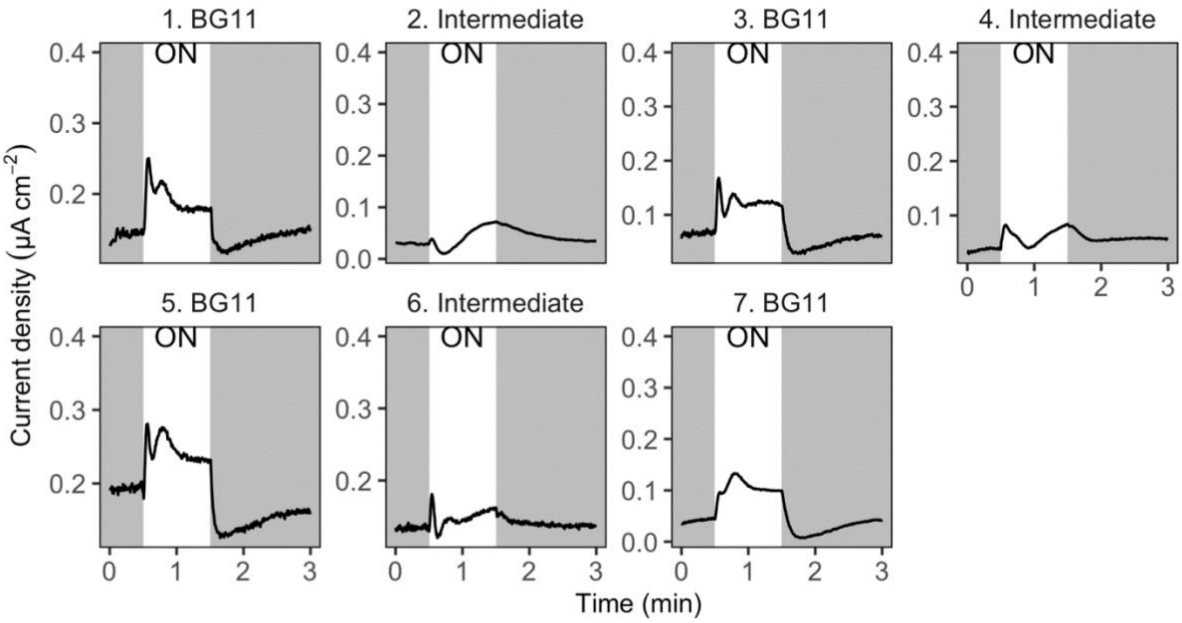


**Supplementary Figure 12** Photocurrent profiles of whole wild-type cells transferred into different electrolytes. Number indicates the order in which the chronoamperometry experiments were performed. Electrolytes tested were: (1,3,5,7) BG11 medium (pH 8.5); (2) a 1/10 dilution of spheroplast buffer (10 mM HEPES buffer with 10 mM $MgCl_2$, 5 mM sodium phosphate and 0.5 M sorbitol (pH 7.5)) in water, which has the components of spheroplast buffer and the osmolarity of BG11 medium; (4) BG11 medium with 10 mM HEPES (pH 8.5), which has the pH of original BG11 medium and the buffering capacity of spheroplast buffer; and (6) BG11 medium with 10 mM HEPES (pH 7.5), which has the pH and buffering capacity of spheroplast buffer. Chronoamperometry measurements were recorded at an applied potential of 0.3 V vs. SHE, and under atmospheric conditions at 25 °C. Light conditions used: λ = 680 nm, 50 µmol photons $m^{-2}$ $s^{-1}$ (approximately 1 mW $cm^{-2}$ equivalent). ON, light on. Data one biological sample, presented as the mean of three consecutive light/dark cycles.

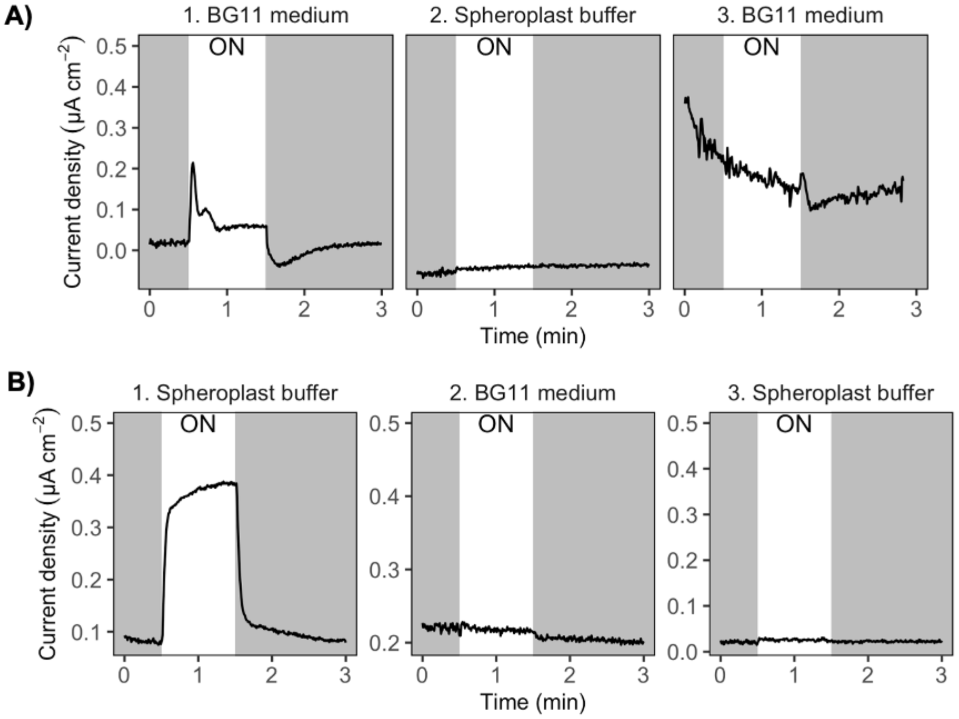


**Supplementary Figure 13** Photocurrent profile in different electrolytes of **A)** whole wild-type cells and **B)** spheroplasts. Number indicates the order in which the chronoamperometry experiments were performed. Chronoamperometry measurements were recorded at an applied potential of 0.3 V vs. SHE, and under atmospheric conditions at 25 °C. Light conditions used: λ = 680 nm, 50 µmol photons $m^{-2}$ $s^{-1}$ (approximately 1 mW $cm^{-2}$ equivalent. ON, light on. Data one biological sample, presented as the mean of three consecutive light/dark cycles

**Supplementary Table 3** Primers used in this study for confirmation of the pilus mutants.

| **Primer** | **Sequence (5' to 3')** |
|---|---|
| pilA1_screen_F | ATAACCAACTAAATCTCTGG |
| pilA1_screen_R | GGTAAGTTACAGCTAGAAGG |
| pilA9-slr2019_screen_F | AATTTGACTAGCAATAGTCC |
| pilA9-slr2019_screen_R | GACCAACATACTGTAACTGC |
| pilT1_screen_F | AAGAATAGTACCGAATTAGC |
| pilT1_screen_R | ATGTTAATCGGTATATTTCC |
| pilB1_screen_F | GGACAGTGGAATGTCCCCCA |
| pilB1_screen_R | GTGGTTCAATGTCGGCAAAA |

**Supplementary Table 4** Combination of primers and their expected sizes for confirmation of the pilus mutants.

| **Strain tested + primers used** | **Expected size (bp)** |
|---|---|
| WT + ΔpilA1 primers | 1481 |
| ΔpilA1 + ΔpilA1 primers | 1724 |
| WT + ΔpilA9-slr2019 primers | 6562 |
| ΔpilA9-slr2019 + ΔpilA9-slr2019 primers | 1821 |
| WT + ΔpilT1 primers | 1700 |
| ΔpilT1 + ΔpilT1 primers | 1521 |
| WT + ΔpilB1 primers | 2654 |
| ΔpilB1 + ΔpilB1 primers | 2084 |

**Supplementary Table 5** Primers used in this study for generation and confirmation of the surface-layer mutant.

| **Primer** | **Sequence (5' to 3')** |
|---|---|
| Slayer_screen_F | GTCACGGCTACCGATGATCT |
| Slayer_ screen_R | CCGAGGCTGTTATCGTCAAT |
| Sll1951leftfor | GATCGAATTCGTCTTTGCCAAGCTGGAATC |
| Sll1951leftrev | GATCGGATCCGAAATTCGGGTCGTTCAAGA |
| Sll1951rightfor | GATCGGATCCGGGAAAATGGTATTGATG |
| Sll1951rightrev | GATCTCTAGATAACTTGATCGCTGGTGCTG |
